# The interplay of membrane tension and FtsZ filament condensation on the initiation and progression of cell division in *B. subtilis*

**DOI:** 10.1101/2025.05.18.654715

**Authors:** Diego A. Ramirez-Diaz, Lei Yin, Daniela Albanesi, Jenny Zheng, Diego de Mendoza, Ethan C. Garner

**Affiliations:** Department of Molecular and Cellular Biology, Harvard University, Cambridge, USA; Instituto de Biología Molecular y Celular de Rosario (IBR), CONICET, Facultad de Ciencias Bioquímicas y Farmacéuticas, Universidad Nacional de Rosario, Rosario, Argentina

## Abstract

The first step of cell division is deforming the planar cell membrane inward towards the cytoplasm. As deforming membranes is energetically costly, biology has developed various protein systems to accomplish this task. The mechanisms providing the force to deform bacterial membranes to initiate division remain unknown. *In vivo* studies have shown the condensation of FtsZ filaments into a sharp ring is required to initiate cell division, an observation mirrored *in vitro* with FtsZ filaments encapsulated inside liposomes. Similarly, the force for membrane deformation in many eukaryotic deforming systems arises from the local crowding of proteins on the membrane surface. As any membrane deforming system works against the membrane tension, here we modulated the amount of lipid synthesis and thus membrane tension in *Bacillus subtilis* to examine: 1) if the condensation of FtsZ filaments by FtsZ bundling proteins serves to overcome the cellular membrane tension to deform the membrane inward and 2) how changes to the membrane tension affect the subsequent invagination of the septum. First, we developed methods to simultaneously measure and modulate membrane tension in live cells. Next, we determined how altering the membrane tension affected the cell’s ability to initiate division with reduced levels of FtsZ bundling proteins. While cells depleted of 2 FtsZ bundling proteins were unable to divide, reducing membrane tension to a given threshold restored their ability to initiate division. Likewise, cells with intermediate levels of FtsZ bundling proteins required a lesser decrease in membrane tension to initiate division. We also found that reductions in membrane tension increase the rate of Z ring constriction, with the constriction rate scaling linearly with the membrane tension. Interestingly, while the constriction rate in wild-type *B. subtilis* is limited by FtsZ treadmilling, the rate of constriction becomes independent of FtsZ’s treadmilling rate when membrane tension is reduced. These experiments give two major insights: First, the filament condensation caused by FtsZ bundling proteins works to overcome membrane tension and deform the membrane inward to initiate division. Second, the rate of septal constriction is limited by membrane tension, suggesting that membrane fluctuations at the tip of the growing septa limit the rate of cell wall synthesis. Finally, our measurements allow the estimation of several physical values of cell division, such as the force required to bend the membrane, but also that the cell membrane provides only 0.1%, a small amount of surface tension relative to the entire cell envelope, indicating 99.9% of the pressure drop occurs across the cell wall. These calculations also indicate that cell division occurs via comparatively very small membrane tension fluctuations relative to the high turgor pressure that exists across the entire cell envelope.

## Introduction

The ability of cells to divide into two is an essential process for all organisms, and accordingly, every cell examined so far encodes a machinery to facilitate its division. At the physical level, any cell division apparatus must first exert enough force to bend the membrane inward against the outward pressure exerted by cellular turgor. While the actomyosin ring is known to generate the constricting force for division in eukaryotic cells, the force-generating mechanisms within the bacterial division machinery that deform the membrane to cell initiate division remain elusive. In bacteria with a cell wall, the division machinery contains two enzymes that polymerize glycan strands and crosslink strands into the existing cell wall. These enzymes associate with other outward-facing membrane proteins thought to regulate their activity, forming what has been termed the “Divisome Core Complex” (Käshammer et al., 2023). The spatial activity of this complex is controlled by two cytoplasmic polymers: FtsA, an actin homolog that binds to the membrane, and the tubulin homolog, FtsZ, which binds to FtsA. There are also FtsZ Bundling Proteins (henceforth referred to as ZBPs), such as SepF, EzrA, and ZapA, which cause FtsZ filaments to laterally associate with each other and can also serve as alternative membrane anchors for FtsZ (Duman et al., 2013; Levin et al., 1999; Pichoff and Lutkenhaus, 2002). These components synergize to divide the cell: FtsA and FtsZ polymerize on the membrane into loose spirals at the future division site and then condense into a sharp “Z ring” by the actions of the ZBPs (Adams and Errington, 2009; Buss et al., 2013; Caldas et al., 2019; Dajkovic et al., 2010; Squyres et al., 2021). Associated with FtsA/FtsZ filaments, the Divisome Core Complex adds new material into the cell wall both before and after FtsZ filaments condense into a sharp Z ring (Squyres et al., 2021). Following FtsZ filament condensation, the Z ring constricts while the enzymes on the opposing membrane surface synthesize the material needed to divide the cell. Dynamic assays have given other insights into the division machinery: 1) FtsZ/FtsA filaments treadmill around the division plane (Bisson-Filho et al., 2017; Li et al., 2018; Perez et al., 2019; Yang et al., 2017). 2) All proteins in the “division core complex” move directionally around the ring as a single unit at a rate limited by - but independent of - FtsZ treadmilling (Bisson-Filho et al., 2017; Lyu et al., 2022; McCausland et al., 2021; Schäper et al., 2024; Squyres et al., 2021; Whitley et al., 2024; Yang et al., 2021), and 3) in *Bacillus subtilis,* the rate of Z ring constriction is not coupled to, but rather limited by the rate of FtsZ treadmilling, *i.e*., increasing or decreasing the treadmilling rate changes the rate of constriction correspondingly (Bisson-Filho et al., 2017; Whitley et al., 2024).

Cell division in *B. subtilis* occurs in 2 phases: *The first phase,* when cell division is initiated, requires FtsZ filaments to condense together to form a sharp Z ring. Z ring condensation requires 1) FtsZ treadmilling (Whitley et al., 2021) and 2) at least 2 out of the 3 ZBPs (Squyres et al., 2021). If FtsZ filaments cannot condense into a ring, Z rings never constrict and cells cannot divide. Notably, the FtsZ-associated cell wall synthetic enzymes remain active and synthesize new cell material at the decondensed rings, but their synthesis does not cause any inward growth of the wall (Squyres et al., 2021). In the *second phase* of division, the Z ring constricts but can only do so after the Z ring has condensed (Whitley et al., 2021). In *B. subtilis*, the Z ring constriction is limited by - and scales with - the rate of FtsZ treadmilling, with slower treadmilling leading to slower constriction and *vice versa* (Bisson-Filho et al., 2017; Whitley et al., 2021). Notably, condensed Z rings (but not uncondensed rings) can constrict even when FtsZ treadmilling is completely inhibited, albeit at extremely slow rates (Whitley et al., 2021).

While we are gaining increasing knowledge about each component of the division machinery’s dynamics, regulation, and functions, one of the most fundamental questions remains unresolved: how the division machinery exerts the force to divide the cell, bending the membrane inward against the cellular turgor. Some models suggest FtsZ’s GTP hydrolysis could generate an inward force, either by inducing FtsZ filaments to convert between straight and inwardly curved conformations (Allard and Cytrynbaum, 2009; Chen and Erickson, 2011; Erickson et al., 1996a, 1996b, 2010; Erickson and Osawa, 2017a; Ghosh and Sain, 2008; Li et al., 2013, 2007; Lu et al., 2000) or by facilitating FtsZ dynamics so that filaments slide past each other to make a constricting coil (Lan et al., 2009). However, in both *S. aureus* and *B. subtilis,* Z rings can still constrict *in vivo* when FtsZ cannot hydrolyze GTP, but only after they have condensed (Monteiro et al., 2018; Whitley et al., 2021). Likewise, purified FtsZ has been shown to inwardly deform liposomes in the absence of GTP hydrolysis *in vitro* (Ent et al., 2014; Osawa et al., 2008). Another model proposes that cell wall synthesis could be the force-exerting process that deforms the membrane (Coltharp et al., 2016; Coltharp and Xiao, 2017), but many organisms like mycoplasma use FtsZ to divide despite having no cell wall or synthesis machinery (Alarcón et al., 2007; Pende et al., 2021; Vedyaykin et al., 2017). Together, these observations suggest there may be some other force-generating process(es) that contribute to the initial membrane bending and subsequent invagination. Notably, a recent model proposed that the periplasm of Gram-positive bacteria could be isosmotic with the cytoplasm, which would reduce (or eliminate) the amount of force needed for both the bending of the membrane and the inward progression of the septum (Erickson, 2017).

The division machinery must conduct the mechanical task of inwardly deforming the membrane, working to overcome the membrane’s rigidity. All membranes have a native membrane tension (Helfrich, 1973) defined as their resistance to deformation. As the membrane tension increases with the difference in osmolarity on either side (turgor pressure), so does the energy needed to deform them. Deforming membranes is energetically costly; deforming membranes of non-walled eukaryotic cells, which are believed to be under less internal pressure than bacteria, requires ∼15kbT of energy (Bloom et al., 1991; Park et al., 2010; Song and Waugh, 1990). To overcome this energetic barrier, eukaryotic cells have evolved a series of systems that inwardly deform their membranes, many of which function by crowding proteins into one location on the membrane surface. These crowding-based systems have been shown to exert membrane bending effects due to 1) the local crowding of proteins generating an entropically driven bending force caused by steric collisions between proteins, generating pressure on one side of the membrane (Busch et al., 2015; Derganc and Čopič, 2016; Hiergeist and Lipowsky, 1996; Lipowsky, 1995; Snead et al., 2019, 2017; Stachowiak et al., 2012, 2010; Steinkühler et al., 2020; Yuan et al., 2023) and 2) amphipathic helices (such as the membrane-associated helix of FtsA) inserting between lipid headgroups causing them to bend apart (Drin and Antonny, 2010; Jarsch et al., 2016; Zimmerberg and Kozlov, 2006), thereby inwardly deforming membranes when crowded into one location. Clathrin deforms membranes by a similar mechanism, but uses the self-assembly of structured proteins to create a defined curvature (Kirchhausen, 2012).

Several *in vitro* studies have demonstrated the condensation of FtsZ filaments can drive membrane deformations, even when FtsZ cannot hydrolyze GTP: 1) When FtsZ-YFP-mts was encapsulated within liposome tubes or giant liposomes, the FtsZ filaments (assembled with either GTP or GMPCPP) induced inward deformations, but only once FtsZ filaments coalesced (Osawa et al., 2008). 2) Likewise, Cryo-electron microscopy (Cryo-EM) of FtsA/FtsZ copolymers within liposomes showed constrictions only where the FtsA/FtsZ filaments were laterally associated and occurred both with GTP and non-hydrolyzable GTP analogs (Ent et al., 2014). Notably, these *in vitro* deformations were observed within liposomes with far lower membrane tension than what exists inside bacteria.

As the division machinery deforms the membrane, it works against the cellular membrane tension, and accordingly, *E. coli* are unable to divide if their membrane is too stiff (Salinas-Almaguer et al., 2022). Membrane tension and membrane surface area are interrelated (Gauthier et al., 2012; Rangamani, 2022). As membrane levels increase around a cell surface, membrane tension decreases, as the increased surface area to volume creates a “hidden membrane reservoir”, where readily available excess membrane is stored in the form of thermal membrane fluctuations **(Fig. 1A)** (Raucher and Sheetz, 1999). This membrane reservoir stored within the membrane fluctuations can exist without the membranes showing visible invaginations or tubes (Ayala et al., 2023). In other words, all membranes undergo thermal fluctuations, and increasing the amount of membrane causes a higher amount of fluctuations, creating a lower membrane tension. As we can modulate the ability of FtsZ filaments to condense by controlling the expression of FtsZ bundling proteins, we worked to examine the interplay of membrane tension and FtsZ bundling proteins on the initiation and progression of cell division in *B. subtilis*.

**Figure 1.**
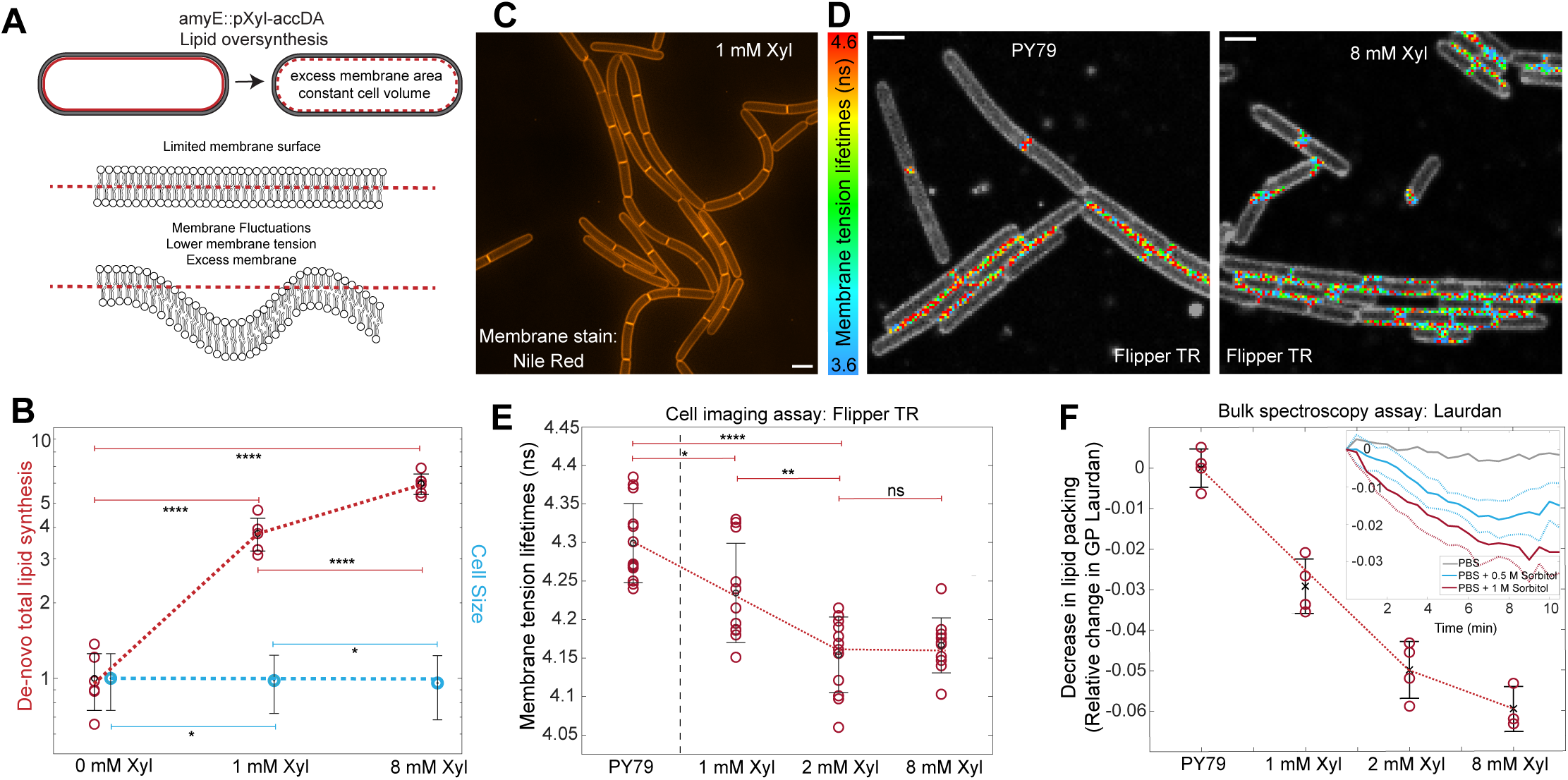
Modulation and measurement of membrane tension *in vivo.* All error bars are SD. **A.** B. Schematic of the strategy used to generate *B. subtilis* cells with membrane excess (reduced membrane tension) by titrating the overexpression of *accDA* to increase phospholipid synthesis. **B and C. B)** *De novo* total lipid synthesis was quantitated by measuring the incorporation of [^14^C] acetate into bSW305 (*amyE::tet-pXyl-accDA)* cells. At a low induction (1 mM xylose), cells produced approximately 4-fold more lipids with no change in **C)** cell morphology as observed by SIM microscopy of cell membranes stained with Nile Red. P-values are in Table S1. Scale bar in C is 1 µm. **D.** Fluorescence lifetime microscopy images of cells stained with Flipper-TR dye. A color code per pixel reflects the results of the double exponential fit to the fluorescent lifetimes: red pixels indicate high membrane tension and blue low membrane tension. Scale bar is 1 µm. **E.** Plot of FlipperTR lifetimes at different *accDA* inductions in bSW305. Photons from different single cells were analyzed. Each data point corresponds to the fitted value gained from all the cells within one field of view. P-values are in Table S1. **F.** Change in the generalized polarization of Laurdan dye of cells with different inductions of *accDA*. **Inset -** Change in lipid packing over time when cells were subjected to hyperosmotic shocks.

### Modulating and measuring membrane tension

To explore how membrane tension affects cell division, we needed means to 1) measure and 2) modulate membrane tension in live *B. subtilis* cells. First, we required a way to incrementally reduce the membrane tension in *B. subtilis* by increasing the rate of cellular phospholipid synthesis, thereby increasing the amount of excess membrane **(Fig. 1A)**. In other words, by inducing an excess of phospholipids, we aimed to generate *B. subtilis* cells with a reduced membrane tension and a reservoir of membrane in the form of membrane fluctuations. For this, we created a strain (bSW305) containing a second copy of *accDA,* an upstream gene in phospholipid synthesis (Kitahara et al., 2022), under xylose inducible control. *accDA* overexpression was previously shown to increase the total amount of phospholipids per cell (Mercier et al., 2013). At low *accDA* inductions (1 mM and 2 mM), the rate of cell growth was comparable to wild type, but higher inductions (8 mM xylose) appeared to be detrimental in stationary phase **(Fig. 1SA)**. We then quantified the de novo total lipid synthesis rate as a function of xylose inductions and the cell size. At 1 mM xylose, lipid synthesis increased by ∼4-fold **(Fig. 1B)**, and cells showed no major changes in their morphology or size relative to wild type when imaged with fluorescence microscopy **(Fig. 1C**). Cell size, cell length, and overall appearance were not highly affected at 2 mM xylose inductions **(Fig. S1B, S1C)**. However, at 8mM xylose, where there was a ∼6-fold increase in phospholipid synthesis **(Fig. 1B)**, cells contained internal lipid clusters near the tips of cells **(Fig. S1D)**.

Next, we sought to measure the membrane tension inside *B subtilis* cells. For this, we used the Flipper TR dye, which changes its fluorescence lifetime with respect to membrane tension **(Fig. 1D)**. Flipper TR has been previously used to measure membrane tension in different organisms and reductions in membrane tension following hyperosmotic shocks in both eukaryotic cells and vesicles (Colom et al., 2018; Roffay et al., 2023). We first validated this assay *in vitro*, where our measurements of membrane tension of GUVs with different lipid compositions using Flipper TR agreed with previous reports (Colom et al., 2018; Roffay et al., 2023) **(Fig. S1E, S1F)**. Next, we acquired FLIM measurements of cells in different fields of view for both wild-type cells (PY79) and our *accDA* overexpressing strain induced at different xylose levels. Relative to the membrane tension in wild-type cells, the membrane tension decreased when *accDA* was induced with 1 mM and 2 mM xylose (∼0.065 and 0.145, respectively), but there was no further decrease at higher inductions **(Fig. 1E)**. This demonstrated that, as expected, the plasma membrane can incorporate a certain amount of phospholipids into the membrane reservoir, but once its capacity is exceeded, the tension no longer decreases with increased membrane, as the excess membrane begins forming multilamellar vesicles (membrane folds inside the cell), as seen in **Fig S1D**.

To validate the Flipper TR observed changes in membrane tension, we conducted bulk fluorescence spectroscopy assays in *B. subtilis* cells using Laurdan, which interdigitates between lipids and reports on lipid packing by changing its emission spectrum (Bagatolli, 2012; Scheinpflug et al., 2016). Using the generalized polarization coefficient, we assayed lipid packing as a function of *accDA* induction. Similar to our Flipper TR results, we observed a consistent decrease in generalized polarization (GP) **(Fig. 1F)** between *accDA* inductions of 1 to 2 mM xylose, with a smaller decrease occurring between 2 and 8 mM xylose. These decreases in lipid packing as a function of membrane excess, coupled with the fact that the cells were roughly the same size, indicate that the membrane surface area per cell does indeed increase with *accDA* induction. As a further verification, we subjected wild-type cells to hyperosmotic shocks while measuring changes in lipid packing with Laurdan. Hyperosmotic shocks reduced GP proportional to the strength of the osmotic difference **(Fig. 1F-insert)**: with an osmotic gradient change of 1M sorbitol, there was a maximal change in GP (∼0.03), a value comparable to cells with *accDA* induced with 1 mM xylose. These experiments demonstrate that increased *accDA* induction increases the membrane surface area to volume ratio inside *B. subtilis,* thereby reducing the membrane tension.

To verify the *accDA*-induced decreases in membrane tension did not arise from changes in the cell’s lipid composition, we characterized the polar headgroups profile with thin-layer chromatography, finding that excess membrane caused no major changes in PE, PG, or cardiolipin content **(Fig. S1G)**. Next, we assessed whether *accDA* overexpression caused major changes in the cell’s lipid profile using mass spectrometry. The membrane fluidity of *B. subtilis* is commonly characterized by the ratio of iso+linear to ante-iso fatty acids: iso+linear fatty acids promote high lipid packing while ante-iso fatty acids reduce packing and increase membrane fluidity (Diomandé et al., 2015; Mercier et al., 2012). We measured the iso+linear/ante-iso fatty acid ratio in cells induced with 0 and 8 mM xylose. At 8mM xylose, cells had a small increase in their iso+linear/ante-iso fatty acid ratio **(Fig. S1H)**. As an increase in iso+linear/ante-iso fatty acid ratio should cause a higher membrane packing (increase in GP Laurdan) and an increase in membrane tension (increase in Flipper TR lifetimes), the decrease in membrane tension and lipid packing observed in the presence of excess membrane cannot be explained by changes to the fatty acids profile.

### Membrane tension and FtsZ condensation effects on division initiation

With our assays established, we worked to examine the interplay and effects between the opposing forces of membrane tension and the ZBP-driven condensation of FtsZ filaments on cell division. Previous *in vitro* studies have shown that FtsZ filament condensation can induce membrane deformations and constriction necks in GUVs under low membrane tension (Ramirez-Diaz et al., 2021) **(Fig. 2A)**, but critically, this deforming process depends on - and works against - the membrane tension. For this effort, we created a *B. subtilis* strain (bDR026) with a second copy of *accDA* under an IPTG-controlled promoter, which is much easier to titrate precisely than the xylose promoter. To assay what *accDA* inductions did not exceed the capacity of the membrane reservoir, we titrated *accDA* expression and examined the cell membranes with electron microscopy and microscopy of fluorescently stained membranes **(Fig. S2A, S2B).** We did not observe any membrane invaginations or vesicles across 0 – 25 µM IPTG, but we did observe abnormal invaginating lipid structures when inductions exceeded 30 µM, indicating inductions beneath 25 µM were only adding lipids into the membrane reservoir. We used this IPTG *accDA* inducible construct to next engineer a *B. subtilis* strain (bDR112) where we could also control FtsZ condensation (having *ezrA* expression under Xylose control at its native locus in a *ΔsepF* background), and with an mNeonGreen fusion to FtsA at the native locus. We also made another strain (bDR110) identical to bDR112 that lacked the mNeonGreen fusion to FtsA. Henceforth, we will refer to *ΔsepF* cells depleted of EzrA as “Z Bundling Protein deficient” (or “ZBP deficient”). Similar to our previous observations (Squyres et al., 2021), when EzrA was depleted in *ΔsepF* cells, FtsZ filaments were unable to condense, and cells could not initiate division, resulting in long cells **(Fig. 2B)**. Furthermore, when *ezrA* was induced with 1mM Xylose, FtsZ filaments condensed, and cells subsequently divided, yielding a length distribution similar to wild type **(Fig. 2D)**. Remarkably, when we expressed excess *accDA* using 25 µM IPTG in EzrA depleted *ΔsepF* cells, these ZBP deficient cells regained the ability to divide **(Fig. 2C)**. Correspondingly, ZBP deficient cells were extremely long at inductions beneath 15 µM IPTG, but at IPTG inductions at or beyond 15µM IPTG cells gradually became smaller, indicating the presence of cell division events – 15µM IPTG being the transition point allowing division **(Fig. 2D, Fig S2B)**. At 25 µM IPTG, cell length distribution became similar to cells with *ezrA* induced with 1 mM xylose. Accordingly, the number of long cells (> 10µm) decreased as a function of membrane excess, with 87% at 5 µM IPTG and only 21% at 25 µM IPTG **(Fig. 2D - inset)**. Thus, reducing membrane tension allows cells with insufficient FtsZ bundling proteins to initiate cell division. The fact that FtsZ filaments condense into rings after they deform membranes is not surprising given that 1) FtsZ filaments have a weak affinity to laterally associate (Bramhill and Thompson, 1994; Erickson et al., 1996b; Guan et al., 2018; Szwedziak et al., 2014), 2) cells still contain the FtsZ bundling protein ZapA, and 3) inwardly curved filaments, such as MreB, FtsA and FtsZ (Erickson and Osawa, 2017b; Khanna et al., 2021; Nierhaus et al., 2022; Salje et al., 2011) are attracted to negative Gaussian membrane curvature (Billings et al., 2014; Hussain et al., 2018; Ursell et al., 2017; Wong et al., 2019), a geometry that is maximal in the cell at the saddle shape of septal invaginations.

**Figure 2.**
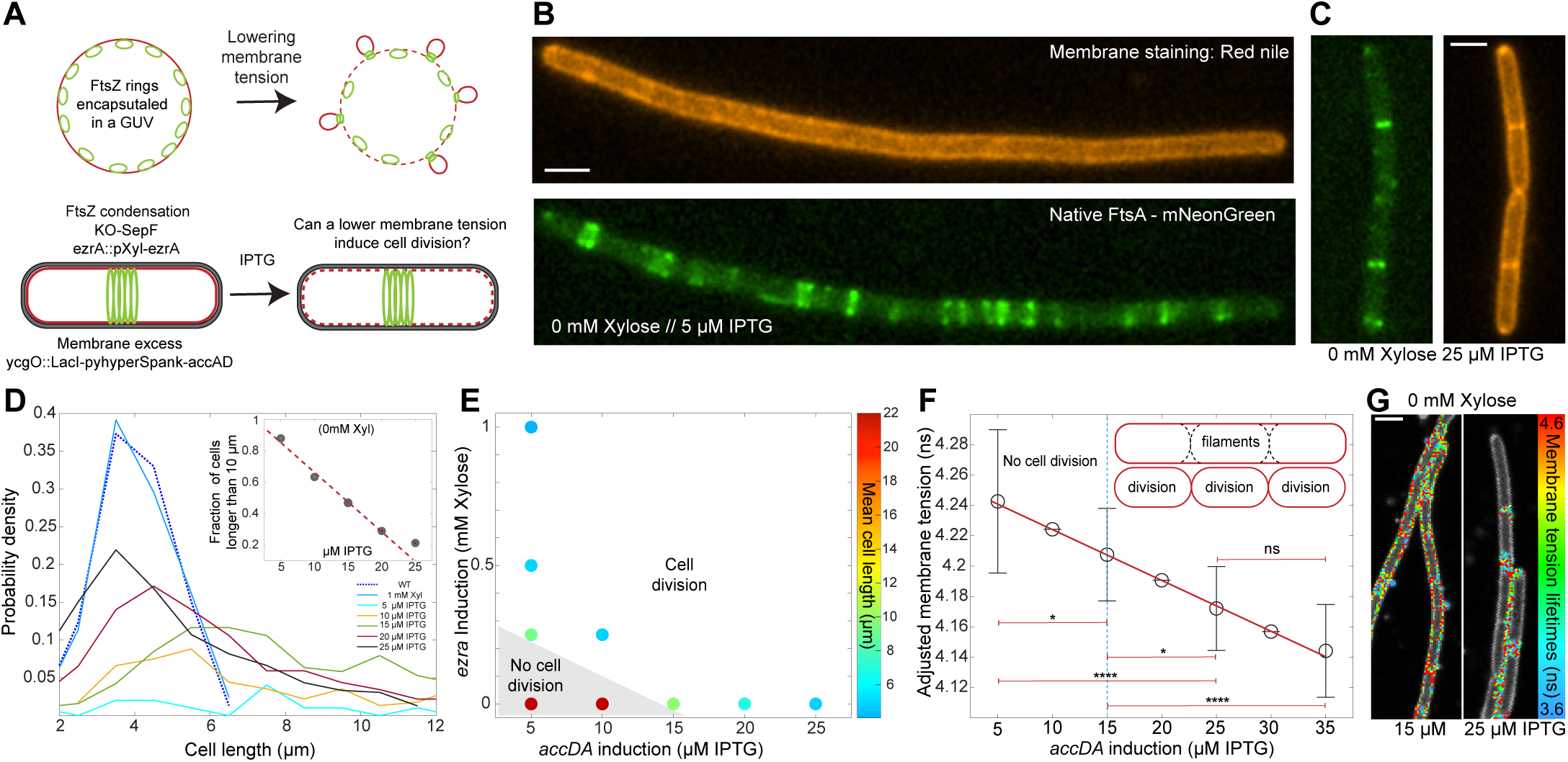
FtsZ condensation by FtsZ bundling proteins serves to overcome membrane tension to initiate division. All error bars are SD. **A.** Schematic of experimental rationale. *Top -* FtsZ inside GUVs are known to deform the membrane when the GUV is beneath a given membrane tension. *Bottom -* Similarly, if the condensation of FtsZ filaments by FtsZ bundling proteins helps to overcome the cellular membrane tension to bend the membrane inward, cells lacking FtsZ bundling proteins should only be able to initiate division at lower membrane tensions. We made 2 *B. subtilis* strains (bDR110 and bDR112) where we could titrate expression of 1) The FtsZ bundling protein EzrA with xylose (regulating filament condensation) and 2) *accDA* with IPTG (regulating the amount of membrane excess) in *ΔsepF* backgrounds. bDR112 also contains mNeonGreen-FtsA(sw) expressed at the native locus. **B.** Long filamented cells (> 10 μm) lacking invaginations formed when bDR112 cells (lacking *sepF*) were depleted of EzrA at low *accDA* induction (5 μM IPTG), and FtsZ only formed decondensed Z rings. Nile Red membrane stain is orange, mNeonGreen-FtsZ is green. EzrA was depleted by growth in 0 mM xylose for 4 hours. **C.** Smaller cells (∼4 μm) capable of dividing formed when *accDA* was induced at higher levels (25 μM IPTG) in bDR112 cells depleted of EzrA. Scale bar is 1 µm. **D.** Cell length distributions as a function of *accDA* induction in EzrA depleted bDR112 cells. At low *accDA* inductions, the distributions are dominated by long cells (> 10 μm), as in Fig. 2B. At inductions of 15 μM IPTG and higher, cell division is evidenced by the increasing number of smaller cells (emerging peaks around 3-6 μm). **Inset -** The corresponding decreasing fraction of long cells as *accDA* induction is increased. **E.** A phase diagram of cell division plotted as a function of FtsZ filament condensing activity (*ezrA* induction) vs. membrane excess (*accDA* induction). Average cell length for each induction pair is represented by spot color. For these Δ*sepF* cells to divide, they require i) higher *accDA* inductions when EzrA is depleted or ii) higher *ezrA* expression (0.25 mM xylose) at low *accDA* inductions (5 µM IPTG). At the threshold points of cell division along each axis (0.25 mM xylose with 5 μM IPTG, and 15 μM IPTG with 0 mM xylose), cells appear as a mix of small cells and long filaments. Cells were unable to divide when *ezrA* was induced with 0.25 mM xylose and *accDA* with 5 µM IPTG, but they regained the ability to divide with near wild-type lengths when *accDA* induction was increased to 10 µM IPTG, demonstrating filament condensation caused by FtsZ bundling proteins serves to overcome membrane tension to deform the membrane inward. **F.** Membrane tension in EzrA depleted bDR110 cells as a function of *accDA* induction. Cells induced with 5 μM IPTG did not divide compared to cells induced at 15µM IPTG and above. To correct for the difference in surface area between filaments and dividing cells (“adjusted membrane tension”), we used cells induced with 5 μM IPTG (*accDA)* and 1mM xylose (*ezrA)* as a proxy for the 5 μM IPTG point, as these cells can divide (see main text). Intermediate values (10, 20, 30 μM IPTG) were linearly interpolated. **Inset** - Diagram of the difference in membrane surface area in dividing and filamented cells. P-values are in Table S1. **G.** Fluorescence lifetime microscopy images of EzrA depleted bDR110 cells stained with Flipper-TR membrane tension dye at different *accDA* inductions. Scale bar is 1 µm.

If FtsZ bunding by ZBPs serves to overcome membrane tension, there should be a relationship between the level of bundling proteins (such as EzrA) and the amount of excess membrane (controlled by *accDA* induction) on the ability of cells to divide. By simultaneously modulating *ezrA* and *accDA* expression, we attained a “phase diagram” for cell division using the mean cell length as readout, using 9 µm (in length) as the boundary between dividing and filamented cells **(Fig. 2E)**. At minimal levels of membrane excess (*accDA* induction), cells require more FtsZ bundling activity (EzrA induction) to divide; conversely, cells with less membrane excess required more bundling activity. Furthermore, cells were able to divide at intermediate inductions of *accDA* and *ezrA* but were unable to divide if either (or both) inductions were insufficient: at 0.25 mM xylose *ezrA* induction and 5 µM IPTG *accDA* induction, cells were unable to divide and formed long filaments (mean cell length of 9.25 µm), but this division defect could be rescued by inducing *accDA* with 10 µM IPTG (mean cell length of 4.9 µm). Thus, 1) both membrane excess and FtsZ condensation by bundling proteins affect the cell’s ability to divide, and 2) the amount of bundling proteins required decreases as the amount of membrane excess increases, demonstrating FtsZ filament condensation serves to overcome membrane tension to initiate division.

To examine the relationship between excess membrane and cell division in the context of membrane tension, we measured membrane tension in the above strain across a range of *accDA* inductions in EzrA-depleted *ΔsepF* cells **(Fig. 2F, 2G, S3C)**. Cells with *accDA* induced at 15, 25, and 35 µM IPTG grew as long chains of cells that contained septa, while cells induced at 5 µM formed long cells lacking septa. Notably, cells with *accDA* induced at 5 µM IPTG showed a significant decrease in membrane tension compared to the chained cells containing septa that arose when *ezrA* was induced at 1mM xylose, a condition that does not require excess membrane for division **(Fig. S2C)**. This reduction in tension likely arises because the long cells lacking septa did not require the membrane surface area needed to build the septa **(Fig. 2F - inset)**, thereby resulting in an *accDA*-independent reduction in membrane tension. Thus, to correct for the difference in membrane surface area at 5 µM IPTG (the lowest *accDA* induction for these experiments) where cells have no septa, as a proxy, we used the membrane tension taken from cells with *ezrA* induced at 1 mM xylose, an induction condition in the same strain with the same *accDA* levels where cells can divide. Plotting the resultant tensions revealed a linear (R^2^ = 0.99) decrease in membrane tension as a function of *accDA* induction **(Fig. 2F)**. The decreasing tension across this induction range further demonstrated the excess membrane created by this range of *accDA* inductions was added to the membrane reservoir. Overall, these experiments demonstrate two points. First, in ZBP deficient cells, cells cannot initiate division if their membrane tension is above 4.21 ns (10µM IPTG *accDA* induction) unless their membrane tension is reduced beneath this threshold. Second, the condensation of FtsZ filaments by ZBPs allows the division machinery to overcome high membrane tension and bend the membrane inward to initiate cell division.

### Membrane tension and septal constriction

We next examined the effect of membrane tension on the second step of cell division, the inward progression of the septa that occurs after cell division is initiated. As before, in EzrA depleted (0 mM xylose) *ΔsepF* cells at low *accDA* induction (5 µM IPTG), cells were long with decondensed FtsZ rings and could not divide. Likewise, when cells were *accDA* induced with 25 µM IPTG, FtsZ filaments condensed into a sharp ring and subsequently constricted **(Fig. 3A)**. At *accDA* inductions of 5µM and 10µM IPTG, Z rings remained decondensed with a large width (0.94 µm and 0.7 µm respectfully). At 15 µM IPTG, the induction threshold where cells were able to initiate division, Z rings showed a sharp reduction in their width (0.44µm) and became thinner at each induction beyond 15µm **(Fig. 3A, 3B - inset).**

**Figure 3.**
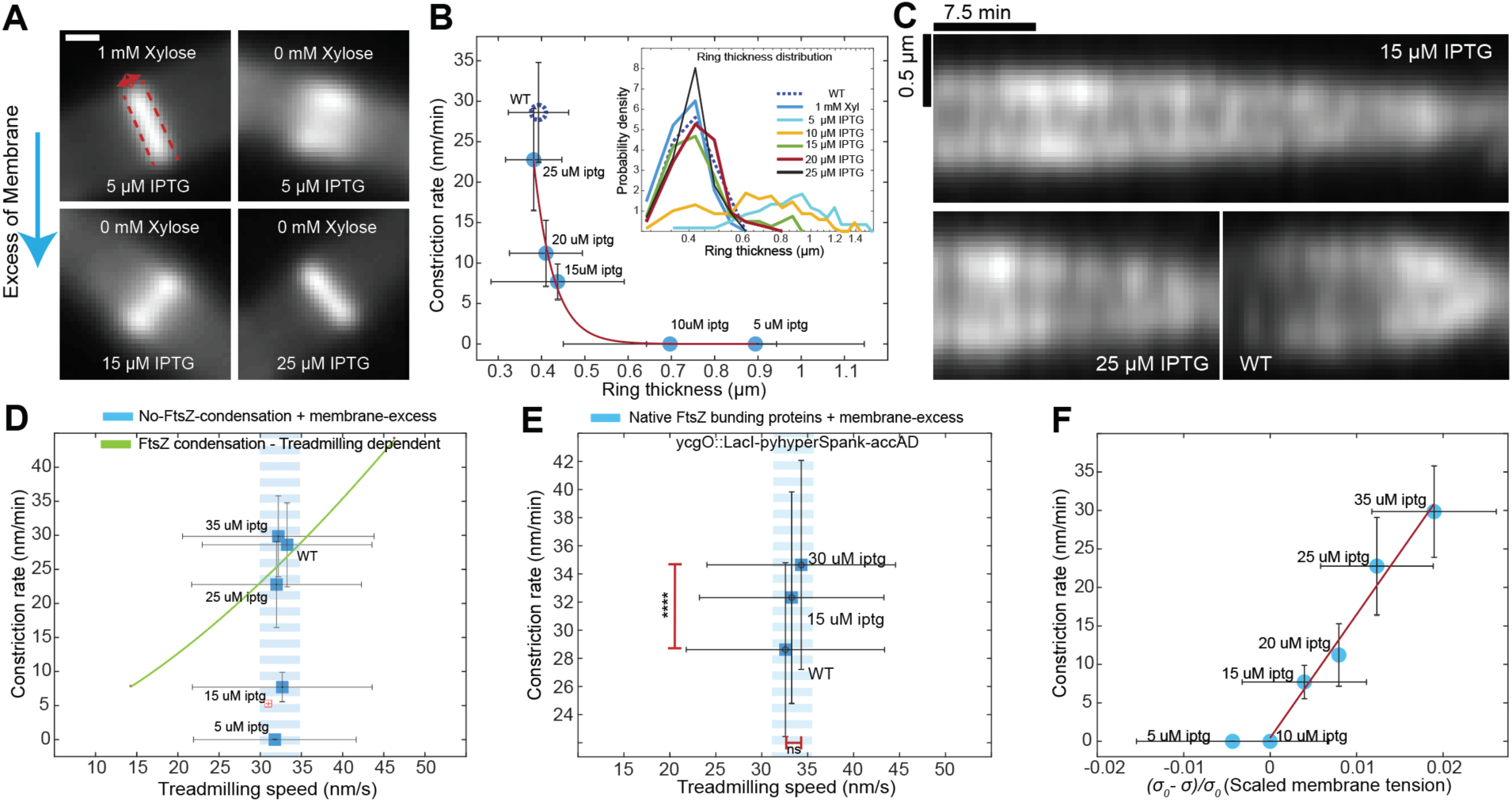
Increasing membrane excess leads to faster Z ring constriction. All error bars are SD. **A.** Time-averaged images of FtsZ rings in *ezrA* depleted bDR112 cells at various *accDA* inductions (IPTG) relative to cells with high *ezrA* induction (1mM xylose). Dotted red line shows an example measurement of ring width. Scale bar is 1 µM. **B.** Plot of Z ring constriction rates vs. ring thickness at different *accDA* inductions in EzrA depleted cells. **Inset** - Thickness distributions of Z rings in bDR112 cells at different *accDA* inductions. Z rings transition from a wide state (∼0.9 μm) at 10 μM IPTG to a compressed state of ∼0.4 μm at 15 μM IPTG. **C.** Kymographs of Z rings showing faster constriction rates at increasing amounts of IPTG controlled *accDA* induction in EzrA depleted bDR112 cells. As before, cell division was only observed at 15 μM IPTG and higher. **D.** FtsZ treadmilling speed vs. Z ring constriction rate at different *accDA* inductions in ZBP deficient cells. While the constriction rate (*blue squares*) increases as membrane tension is reduced in ZBP deficient bDR112 cells, the speed of FtsZ filament treadmilling remains constant. This contrasts with the previously observed relationship (*green line*) in *B. subtilis* cells with wild-type membrane tension, where the treadmilling speed limits the constriction rate (Bisson-Filho et al., 2017). Error bars are SD. **E.** FtsZ treadmilling speed vs. Z ring constriction rates at different *accDA* inductions in cells with native ZBP expression. As in Fig. 3D, constriction rates also increase with *accDA* overexpression in these otherwise wild-type cells. As above, the speed of FtsZ filament treadmilling remains constant across *accDA* inductions. P-values are in Table S1. **F.** Ring constriction rates plotted against scaled membrane tension. Constriction rates scale linearly with membrane tension in ZBP-deficient cells at *accDA* inductions above 15 µM.

To measure Z ring constriction rates at different levels of excess membrane, we constructed kymographs along the division plane and measured their slope across different *accDA* inductions **(Fig. 3C).** Compared to wildtype cells, cells induced with 15 µM IPTG had an ∼4 fold slower constriction rate than wild type, and the constriction rate increased at higher inductions. Plotting the ring constriction rates against their thickness at each *accDA* induction revealed that, once excess membrane exceeded the threshold to initiate division, increasing the excess membrane caused both increasingly thinner rings and faster constriction rates **(Fig. 3B).** The increased constriction rates did not come from thinner Z rings building thinner septa (or *vice versa*), as measuring the septal thickness with TEM (Transmission Electron Microscopy) revealed the septal thickness was unchanged across *accDA* inductions, even though Z rings were wider at lower *accDA* inductions **(Fig. 3A, S3A,B)**. Furthermore, immunoblots revealed that FtsZ levels did not change across the *accDA* induction range used **(Fig. S3C)**.

In rich media, the rate of Z ring constriction in wild-type B. subtilis is limited by the rate of FtsZ treadmilling; faster treadmilling causes faster constriction, and slower treadmilling causes slower constriction (Bisson-Filho et al., 2017; Whitley et al., 2021) **(Fig. 3D**, *green line*). To explore if the increase in constriction velocity caused by increased membrane excess arose from changes in FtsZ’s treadmilling rate, we measured FtsZ’s treadmilling rate at different *accDA* inductions. Surprisingly, FtsZ’s treadmilling speed remained constant across the induction range, with a value very similar to wild-type cells (∼33 nm/s) despite the Z rings constricting at different rates **(Fig. 3D)**. We then tested if increased excess membrane also increased the rate of ring constriction in otherwise wild-type cells (cells with native *ezrA* and *sepF* expression) and its relationship to FtsZ treadmilling. Again, we observed faster constriction rates for *accDA* levels (15 µM IPTG and 30 µM IPTG) while treadmilling rates remained unaffected **(Fig. 3E)**. Together, these experiments reveal that in rich media, the rate at which the Z-ring constricts in *B. subtilis* is limited by the amount of cellular membrane, and this increased constriction did not arise from (nor cause) a change in FtsZ levels, width of the division septa, or FtsZ’s treadmilling rate.

How could an increased amount of membrane cause an increased rate of cell constriction? This could arise from the resultant decrease in membrane tension. Notably, at *accDA* inductions above the 15 µM IPTG division threshold we observed a fairly linear correlation between the constriction rate and membrane tension (R^2^=0.978) **(Fig. 3F)**. The increasing rate of constriction occurring with reductions in membrane tension agrees with a previous observation in *E. coli,* where the rate of Z ring constriction increased when cells were subjected to hyperosmotic shocks (Sun et al., 2021). Thus, the cellular membrane tension somehow limits the rate at which the Z ring constricts in *B. subtills*.

In total, the above experiments reveal the amount of available membrane and the resultant membrane tension limits cell division in two general ways: i) the initiation of cell division, where the membrane is bent inward against tension by filament condensation, and ii) the rate at which the septa is synthesized to divide the cell.

### Physical estimates

By connecting prior *in vitro* evidence with the -here-presented phenomenology, we can estimate certain physical parameters relevant to *B. subtilis* cell division. The first aspect to consider is the FtsZ intrinsic curvature relative to the curvature of the cellular membrane. The curvature of FtsZ filaments depends on the tension of the membrane they bind to: FtsZ-YFP-mts assembled within low tension GUVs forms rings matching the native curvature of FtsZ filament (>1 µM^-1^), but a small increase in membrane tension (∼0.03 ns) **(Fig. S4A)** causes FtsZ to transition into long bundles with lower curvatures matching the GUV diameter (<0.1 µM^-1^). This observation, along with previous studies (Ganzinger et al., 2020; Ramirez-Diaz et al., 2021), demonstrates membrane tension affects whether membrane-associated filaments 1) inwardly bend the membrane so it matches the filament curvature or 2) deform themselves to match the membrane curvature.

Given these findings, we can estimate A) how much membrane excess is needed and B) the depth of the inward membrane required to initiate cell division in Δ*sepF* EzrA depleted cells. As low membrane tension favors FtsZ rings with a smaller diameter (higher curvature), we can use Z ring thickness as an indirect measure of membrane excess in the cellular context. A decrease in Z ring thickness within a cylindrical *B. subtilis* cell could be explained by FtsZ rings inwardly deforming the membrane with a deformation depth *h* **(Fig. 4A)**. As wide Z rings in cells with high membrane tension become gradually thinner as we decrease membrane tension **(Fig. 3A, 3B)**, by denoting the initial surface area of the cell as *A*, the cell length as *L*, and the surface area change Δ*A* due to the excess membrane (aka, the hidden membrane area), the ratio of the amount of excess to initial membrane 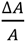 can be inferred by the difference in Z ring thickness Δ*z*. Assigning z as the FtsZ ring thickness as a function of *accDA* induction (IPTG), the inward deformation depth *h* can be calculated 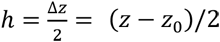 with *z*_0_ being the most relaxed (widest) FtsZ ring thickness occurring at 5 µM IPTG. Given the cylindrical geometry, 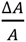 can be expressed as 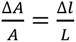 where Δl denotes the “hidden membrane length” stored in the membrane fluctuations, thus 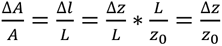 **(Fig. 4C)**. Inducing *accDA* at 15 µM IPTG (the lowest induction at which cell division can initiate) the excess membrane ratio 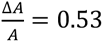 implies that our ZBP deficient cells require 53% more hidden membrane area to initiate cell division. In agreement with this estimation, we measured the excess of membrane synthesis at 15 µM IPTG, finding at this “division threshold,” cells contain 85% more membrane **(Fig S4B)**. Furthermore, in ZBP deficient cells with this amount of excess membrane we can calculate that an inward membrane deformation of *h* ≈100 nm is the minimal threshold to initiate cell division. The validity of this geometric approach arises from the fact that, in the absence of FtsZ bundling proteins, we see minimal lateral interactions between filaments *in vivo* (decondensed rings) that could contribute energetically to deform the membrane.

**Figure 4.**
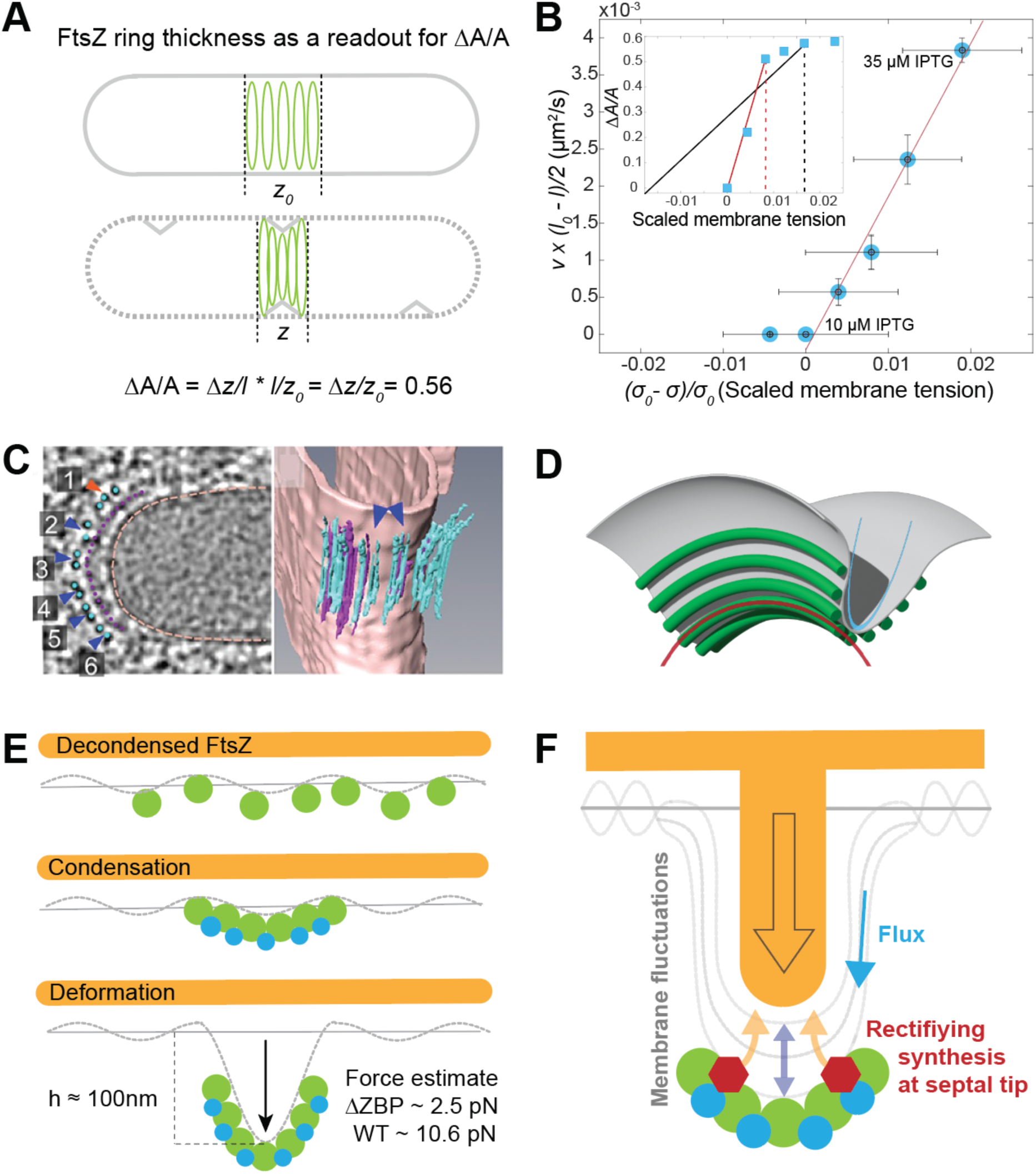
Estimates and Models. A. The excess of membrane 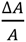 required to initiate cell division in ZBP deficient cells can be estimated by the difference in Z ring thickness Δz. At the threshold *accDA* induction (15 μM IPTG) where cells begin to divide, 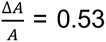, close to our physical measurements (0.85) (**Fig. S4B**). We also estimate that division initiation requires an initial inward membrane deformation of at least *h* = 100 nm. B. By using the constriction rates as proxy for lipid flux velocity *v*, the scaled membrane tension from Flipper TR lifetimes, and the equation 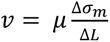, we can estimate membrane tension at different *accDA* inductions: σ_0_ = 0.344 mN/m for cells when *accDA* was induced with 10 µM IPTG. **Inset -** By plotting membrane tension vs excess membrane we see a linear range in the *accDA* inductions (5-15 µM IPTG, *red line*) beneath the threshold where the membrane yielded and cells-initiated division. From this linear range, we estimated the stretch modulus *E* = 5.56 pN mm^-1^ which is equivalent to a force of *f* = 2.54 pN as the force required to initiate division. Lastly, by using the WT interpolated membrane tension (0.350 mN/m), we estimated *E* = 21.08 pN mm^-1^ for wild type cells (*black line*) equivalent to a force of *f_wt_* = 10.57 pN. C. **Tomograms of FtsZ filaments at division septa. Used with permission from** (Khanna et al., 2021) *Left* - Slice through a tomogram of a dividing *B. subtilis* cell. Membrane distal filaments (FtsZ with a Q-rich linker) are blue, membrane-proximal filaments (FtsA) are pink. Blue arrows indicate doublets, and an orange arrow indicates a possible triplet. *Right -* Reconstructed cell membrane, with membrane-proximal and membrane-distal filaments corresponding to the figure on the left. Blue arrows point to doublets of membrane-distal filaments. Similar arrangements were observed in cells containing native FtsZ D. **Model for how FtsZ filaments induce opposing curvatures at the division septa.** After FtsA/FtsZ filament crowding initiates division, the membrane (*grey)* around the septal invagination forms a saddle shape composed of two opposing principle curvatures *(blue and red lines*). FtsA/FtsZ filaments and their condensation may play a role in establishing this geometry: FtsA/FtsZ filaments (*green*) have an intrinsic inward curvature, which may induce (and stabilize) the deformation parallel to the rod width (*red axis*). As expected for a crowding induced deformation, FtsA/FtsZ filaments are spaced around the septal invagination (*blue axis*), an arrangement that induces and stabilizes the membrane deformation along the cell length. E. Model for how the membrane is deformed to initiate cell division. The membrane (dotted line) has a tension that resists deformation. FtsA/FtsZ filaments (green circles) are bound to (and treadmill along) the membrane, but individually cannot deform the membrane. FtsZ bundling proteins (blue circles) cause FtsA/FtsZ filaments to laterally associate. This local filament crowding works to overcome the membrane’s tension to deform it inward, giving the space needed for cell wall synthetic enzymes to begin building the division septa. F. Model for inward progression of the division septa. Membrane fluctuations occur all over the membrane surface as well as at division septa. These fluctuations give the cell wall synthesis enzymes (red hexagons) that are associated with FtsA/FtsZ filaments room to insert new cell wall material at the base of the septa. Each insertion event rectifies the inward membrane fluctuation, thereby causing the division septa to ratchet inward. Membrane flux of into the invagination may also increase membrane fluctuations at the tip of the invagination.

We next examined how membrane tension affected the second step of division, the inward progression of the division septa that occurs after the initial inward deformation. Given the linear correlation we observed between constriction rate and membrane tension in our cells lacking ZBPs **(Fig. 3F)**, the membrane levels (or tension) appear to be limiting the constriction rate. This suggests that division requires - and is limited by - the flux of lipid material into the invaginating septa from the membrane excess stored in the hidden membrane area and/or native lipid synthesis. A lipid membrane responds to a local perturbation (such as a bend) by stretching or lipids flowing toward the perturbation (Supporting Information S1). Forces arising from FtsZ bending the membrane and the reinforcement of this deformation by cell wall synthesis could impose a membrane tension gradient, which consequently generates a lipid flux towards the tip of the septa. Here, we postulate that σ*_m_*, being the measured membrane tension as function of *accDA* expression, represents a resistive component in such a way that higher levels of AccDA decrease membrane tension and reduce the membrane resistance to flow. Then, the increase in lipid flux velocity can be expressed as 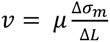 (see derivation in Supporting Information S1), with *μ* being the membrane mobility (Cohen and Shi, 2020; Djakbarova et al., 2021; Shi et al., 2022, 2018), Δσ*_m_* being the membrane tension difference with respect to σ_0_ as the membrane tension at the highest expression level of *accDA* with no cell division and Δ*L* as the cell length difference. This increased lipid flux could explain the faster constriction of Z rings as membrane levels were increased; not only would it provide the lipid needed for the growing septa, but it would also increase the membrane fluctuations at the tip of the septa, which in turn could also the accelerate the rate of inward cell wall synthesis if the addition of new septal material occurs via a Brownian ratchet mechanism.

Accordingly, we can estimate the membrane tension at different *accDA* inductions in physical units (mN/m) using the rate of constriction (*V_c_*) as a function of *accDA* induction as a proxy for the rate of lipid flux (*v*) into the septa. Since we measured the membrane tension σ*_m_* (in ns units) as a function of *accDA* induction, the scaled membrane tension difference is defined as 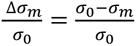 with σ_0_ being the membrane tension at 10 µm IPTG, highest induction where cells no longer initiate division. In addition, we calculated the change in cell length Δ*l* = (*l* − *l*_0_) with (*l*) being the cell length as a function of accDA induction and (*l*_0_) defined as the cell length at the 10 µM IPTG. Using the slope of the plot 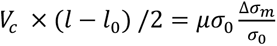 **(Fig. 4B)** and reported values of membrane mobility *μ* ∼ 6 × 10^-4^ μm^3^ pN^-1^s^-1^ (Cohen and Shi, 2020), we can calculate the membrane tension at 10 µM IPTG *accDA* induction as σ_0_ = 0.344 mN/m.

Next, using the membrane tension estimated above, we can estimate the forces that FtsZ condensation exerts on the membrane to initiate cell division in these SepF and EzrA deficient cells. We plotted membrane tension vs membrane excess to estimate the membrane stretch modulus *E* **(Fig. 4B - inset)**. Similar to a typical stress-strain response curve, we observed a linear range (5-15 µM IPTG, red line) of stress before the membrane yielded, a yielding corresponding to the inward membrane deformation that initiates cell division. In this linear range, the stretch modulus is estimated to be *E* = 5.56 pN μm^-1^ **(Fig. 4B – inset***, red line*). For context, lipid membrane tethers can be pulled out of neutrophils and other eukaryotic cells with constant pulling force (Cohen and Shi, 2020), indicating an effective *E*∼0, meaning the cell membrane was not being stretched but rather the hidden membrane area being unfolded. Likewise, our low estimation of *E* indicates that, in our case, the excess membrane is also being unfolded during both the initiation and progression of cell division. Given the difference in FtsZ ring thickness *z* − *z*_0_ = 0.45µ*m* measured between 5-15 µM IPTG, the force required force to inwardly deform the membrane in these ZBP deficient cells containing excess membrane can be estimated as *f* = 2.54 pN. Notably, *in vitro* FtsZ rings within GUVs with low membrane tension exert forces in a similar range (∼1 pN), forces adequate to deform their membranes and generate constriction necks (Ramirez-Diaz et al., 2021). Thus, the force required to initiate cell division in ZBP deficient cells with excess membrane is similar to what is required to deform flaccid GUVs.

We can then also estimate the membrane tension in wild-type cells (PY79) as 0.350 mN/m using the approaches above; from the wild-type FtsZ ring thickness, *z_wt_* − *z*_0_ = 0.5, we calculated that wild-type cells require a change in area 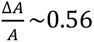 to initiate cell division. Using this value, the stretch modulus for wild-type membranes can be estimated as *E* = 21.08 pN μm^-1^ (**Fig. 4B – inset**, *black line*), and the force required for the membrane deformation needed to initiate cell division to be *f_wt_* = 10.57 pN, 4-fold higher than our ZBP deficient cells with excess membrane, and 10-fold higher than what is required to deform flaccid GUVs.

The above estimates reveal another surprising result, as the turgor pressure in *B. subtilis* has been reported to be ∼10 atm (Whatmore and Reed, 1990), the total surface tension of the entire cell envelope (both the membrane and cell wall) is ∼350 mN/m (using Laplace’s law for a cylinder). This indicates that the cell membrane contributes only ∼0.1% (0.350 mN/m) of the total surface tension strength that maintains cell shape. Importantly, these measurements also reveal that membrane fluctuations (∼0.01mN/m) representing only <0.01% of the total surface tension are sufficient to initiate cell division in wild-type cells. In other words, cell division occurs via comparatively very small membrane tension fluctuations, regardless of the high turgor pressure that exists across the entire cell envelope.

## Discussion

The experimental data above suggest that membrane tension plays a limiting role in both of the two stages of division: the initiation of division and the invagination of the septa.

Regarding the initiation of cell division, our data, along with membrane fluctuation analysis studies in *E. coli* (Salinas-Almaguer et al., 2022), suggest that thermal membrane fluctuations are critical to the initiation of cell division, and increased fluctuations are required in cells with decreased FtsZ bundling proteins. Moreover, we observe a phase space where decreased FtsZ filament crowding on the membrane surface (mediated by FtsZ bundling proteins) can be overcome by a corresponding decrease in membrane tension. This suggests that similar to the many crowding-induced membrane-deforming systems characterized in eukaryotes (Belessiotis-Richards et al., 2022; Boucrot et al., 2012; Brown and Hoh, 1997; Dannhauser and Ungewickell, 2012; Day et al., 2021; Ford et al., 2002; Küey et al., 2022; Kumar et al., 2002; Sargiacomo et al., 1995; Sens and Turner, 2004; Snead et al., 2017; Yu and Schulten, 2013), the local crowding of FtsZ filaments, caused by bundling proteins, may serve to overcome the membrane tension to bend the membrane inward, an energetically costly step in cell division. This model also agrees with past *in vitro* work, including studies finding that 1) FtsZ reconstituted within tubular liposomes only induced observable inward deformations when Z rings coalesced (Osawa et al., 2008) and 2) Cryo-EM work showing liposomes constricted only where there was a high density of FtsA and FtsZ filaments (Szwedziak et al., 2014). Notably, crowding-induced membrane deformations provide a way to exert an inward force on the membrane independent of FtsZ’s GTPase activity, explaining why FtsZ can deform liposomes in the above studies when assembled in the presence of non-hydrolyzable nucleotide. This does not mean this first step of cell division is completely independent of GTP hydrolysis by FtsZ, as treadmilling is required to condense FtsZ filaments (Whitley et al., 2021).

Cryo-EM experiments have shown that FtsZ filaments are spaced equally around the septal invaginations in *B. subtilis*, as expected for membrane deformations caused by entropic repulsion (Khanna et al., 2021; Szwedziak et al., 2014) (**Fig. 4C**). The FtsZ crowding induced deformations, combined with the inward curvature of FtsA/FtsZ filaments, may create the membrane’s saddle shape required for cell division: 1) the inward curvature of FtsA/FtsZ filaments pinches the membrane inward (**Fig. 4D – red lines**) while 2) their condensation creates a bend perpendicular to and in the opposite direction of the first, resulting in the negative gaussian curvature of the membrane at the division site **(Fig. 4D – blue lines).**

Our calculations – based entirely on physical principles – suggest that the membrane only contributes ∼0.1% of the total surface tension of the entire cell envelope (∼350 mN/m), with the pressure drop across the membrane being only a minuscule fraction of that across the cell wall. While this data does refute the model that the periplasm is isosmotic with the cytoplasm, it does agree with the idea that the cell wall, and not the membrane, holds back the vast majority of the cell’s internal pressure in Gram-positive bacteria (Erickson, 2017) where the majority of the turgor might be neutralized by a Donnan potential created by the high density of cations bound by peptidoglycan and teichoic acids in the cell wall (Kern et al., 2010; Thomas and Rice, 2014). Furthermore, our calculations suggest that membrane tension fluctuations (∼0.01mN/m), representing only <0.01% of the total cell surface tension, are sufficient to initiate cell division regardless of turgor.

By conducting physical calculations based on our measurements, we could estimate the forces needed for the inward membrane deformation in different cellular conditions. For cells with reduced membrane tension that lack FtsZ bundling proteins, we estimate the forces to be 2.54 pN, a force similar to measurements of the forces exerted by FtsZ rings (∼ 1 pN) deforming flaccid (low tension) lipid tubes (Ramirez-Diaz et al., 2021), as well as theoretical estimations of this force: 0.35 – 2.45 pN (Osawa and Erickson, 2018). Such forces are sufficient to drive a critical inward deformation of 100 nm, as estimated above, the membrane deformation required to initiate cell division in ZBP deficient cells **(Fig. 4A,E).** Using similar assumptions, we estimate that wild-type cells require a force of ∼10 pN to bend the membrane inward. While our data suggests the local crowding of FtsZ filaments provides the force to deform the membrane inward, membrane synthesis might also play a role as it reduces membrane tension. However, when lipid synthesis is halted in *B. subtilis* cells with cerulenin, the cells continue to grow and divide for ∼1 generation (Pulschen et al., 2016), suggesting wild-type cells contain enough membrane in the reservoir to conduct division without the need for additional lipid synthesis. The forces underlying these membrane deformations could arise from various non-exclusive mechanisms, including i) the local crowding of proteins inducing entropically driven membrane deformations (Busch et al., 2015; Derganc and Čopič, 2016; Hiergeist and Lipowsky, 1996; Lipowsky, 1995; Snead et al., 2019, 2017; Stachowiak et al., 2012, 2010; Steinkühler et al., 2020; Yuan et al., 2023) and ii) the local crowding of amphipathic helices (such as in FtsA) inserting between lipid headgroups on one face of the bilayer (Drin and Antonny, 2010; Jarsch et al., 2016; Pichoff and Lutkenhaus, 2005; Zimmerberg and Kozlov, 2006).

Our data also suggests that during the second stage of cell division, the amount of cellular membrane around the cell is limiting for septal invagination, as constriction accelerates with increasing amounts of membrane. Therefore, as membrane levels increase, so does the velocity of membrane flux into the division site, which correspondingly increases the amount of membrane fluctuations (reduced membrane tension) at the tip of the invaginating septa. We propose that the inward progression of the septa may occur via a Brownian ratchet mechanism, where membrane fluctuations at the growing septal tip create room for enzymes to insert new cell wall. This synthesis would then rectify the membrane fluctuations via mechanical stabilization, causing the septa to gradually move inward with each insertion event **(Fig. 4F)**. This would mean septal constriction is not driven by cell wall synthesis but rather by Brownian fluctuations. Notably, this model agrees with *E. coli* studies that found the rate of Z ring constriction can be accelerated by hyperosmotic shocks (Sun et al., 2021). A synthesis rectified Brownian ratchet may be a common mechanism underlying division in walled organisms, as a study in *Schizosaccharomyces pombe* found the constriction rate 1) increased when turgor was reduced, 2) required the synthesis of glucans, and 3) was independent of the actomyosin contractile ring at later time points, leading the authors to a similar model that division in fission yeast occurs via a rectified Brownian ratchet (Proctor et al., 2012).

Given A) membrane tension is more limiting for constriction than FtsZ treadmilling, but B) treadmilling limits constriction in cells with normal membrane tension, it may be that the inward progression of the septa occurs by a continuous process of FtsZ locally shaping membrane fluctuations at the septa, and these local fluctuations are then rectified by cell wall synthesis. Our two different observed “modes” of cell division, where either 1) membrane tension or 2) FtsZ treadmilling is limiting might explain differences between division in *B. subtilis* and *Staphylococcus aureus*: In contrast to treadmilling limiting Z ring constriction in *B. subtilis* (Bisson-Filho et al., 2017; Whitley et al., 2024), in *S. aureus* the constriction rate is independent of FtsZ’s treadmilling (Schäper et al., 2024), and i may be this discrepancy arises due to a difference in membrane tension in the two organisms.

The role of the local crowding of proteins in deforming membranes is a mechanism implicated in an increasing number of membrane-deforming systems. The role of FtsZ crowding in deforming the membrane may be a deeply rooted evolutionary conserved process: ZapA and SepF are highly conserved within FtsZ-containing bacteria, and most archaea have at least one FtsZ homolog that systematically co-occurs with SepF (Pende et al., 2021). Indeed, phylogenetic work has suggested that SepF and FtsZ were part of the division machinery in the Last Universal Common Ancestor (LUCA), being present before archaea and bacteria diverged (Pende et al., 2021). Furthermore, mycoplasmas lack most of the cell division proteins, containing only FtsZ, FtsA, and the bundling protein SepF (Alarcón et al., 2007; Pelletier et al., 2021). Thus, FtsZ crowding-induced membrane deformations might be an important and deeply conserved process underlying the initiation of cell division in both bacteria and archaea.

## ACKNOWLEDGEMENTS

We thank Adam Cohen, KC Huang, Dan Fletcher, Leendert Hamoen, and Harold Erickson for insightful discussions. D.A was supported by Agencia Nacional de Promoción Científica y Tecnológica (ANPCyT) of Argentina, Grants PICT 2019-2019-00694 and PICT 2021-IA-00310 to D.A.. J.Z. was supported by a grant 203276/l/16/Z from the Wellcome Trust and support from the NSF-Simons Center for Mathematical and Statistical Analysis of Biology at Harvard (award #1764269). Electron Microscopy Imaging, consultation, and services were performed in the HMS Electron Microscopy Facility. We also thank the Harvard Center for Imaging Biology for support with FLIM microscopy.

## MATERIALS AND METHODS

### Culture growth

A. *subtilis* strains (PY79, bSW305, bDR018, bDR026) with wildtype FtsZ bundling proteins (natively expressed *ezrA* and *sepF*) were prepared as described below. Strains were streaked from −80°C freezer stocks onto LB agar plates and grown overnight at 37°C. Single colonies were transferred to liquid cultures in CH medium. Cells were placed on a roller drum grown at 37°C. After cultures reached OD600 ∼ 0.5, one serial dilution (1:10) was grown until OD600 ∼ 0.5. For Δ*sepF* cells with *ezrA* under xylose control and *accDA* under IPTG control, single colonies were streaked from −80°C freezer stocks onto LB agar plates with 5mM xylose. After overnight at 37°C, single colonies were transferred in CH medium with 1 mM xylose and cultured on a roller drum grown at 37°C. After reaching a OD600 ∼ 0.5, 40 µL of liquid culture was diluted in 960 µL of fresh media (1:25 dilution) with or without IPTG. Following this dilution, cells were grown for 4-6 hours depending on IPTG levels. This initial (1:25) dilution and long growth period times allowed more than 5 generations in order to achieve *ezrA* depletion.

### Imaging dish and sample preparation

Dishes with No. 1.5 glass-bottom (MatTek) were cleaned with 2% Hellmanex (Hellma GmbH&Co, Milllheim, FRG) overnight, followed by multiple washes with deionized water, then sonicated with 1M KOH for 30 minutes, followed by multiple washes with water. To prepare the imaging sample, 1 uL of cell culture were placed on the glass surface of the imaging dish and covered by a 1.5% agarose pad made with CH media to create a thin liquid film.

### Quantitative analysis of the de novo lipid synthesis and phospholipids and fatty acids profiling

*B. subtilis* PY79, bSW305 and bDR026 strains were streaked from −80°C freezer stocks onto LB agar plates, and grown overnight at 37°C. The strains were inoculated in liquid LB to an OD600=0.04 in the absence or presence of xylose or IPTG as indicated. Cultures were grown with shaking at 37°C to an OD600 ∼ 0.3 and labeled with 2 µCi.ml-1 [14C]-acetate for 60 min. Cells were collected and total lipids were extracted by the Bligh & Dyer method with minor modifications (Bligh and Dyer, 1959). Briefly, the cells were resuspended in 100 µl of water, subsequently 750 µl of chloroform-methanol (1:2; v/v) were added and the samples were incubated ON at -20°C. Samples were centrifuged for 2 min at 14,000 g. The supernatants were recovered and transferred to a new tube containing 250 µl of chloroform and 250 µl of 1 M KCl. The samples were vortexed and centrifuged to promote phase separation. Each organic phase was transferred to a new tube. The aqueous phase was re-extracted with 250 µl of chloroform and combined with the first extraction. The organic phases were evaporated under nitrogen and resuspended in 100 µl of chloroform. Radioactivity was measured on a scintillation counter. Lipids were separated onto thin-layer silica gel plates (Kieselgel 60; Merck) in two dimensions thin-layer chromatography [TLC] using chloroform-methanol-water (14:6:1, v/v/v) in the first phase followed by separation in a second phase with chloroform-methanol-acetic acid (13:5:2, v/v/v). Plates were exposed to a radioactive storage screen and analyzed using an Amersham Biosciences Molecular Dynamics Typhoon™ FLA 7000 scanner. Lipid species were identified by comparison with migrated phospholipid standards and quantitated using ImageJ software.

In order to analyze the fatty acid (FA) content in the above mentioned growth conditions, total lipids from 50 ml unlabeled cultures were prepared as described and derivatized to methyl-esters (FAMEs) by acid-catalyzed esterification/transesterification (Bernert, 1989). Dried total lipids were resuspended in 2 ml of methanol, 2-5% sulfuric acid was added (10-15 drops of 98% w/w sulfuric acid) and samples were incubated at 80 °C for 2 h. After reaching room temperature, 1,5 ml hexane and 1,5 ml of 5% w/v NaCl were added, samples were vortexed and subsequently centrifuged for phase separation. The organic phase was recovered in a clean tube. The aqueous phase was re-extracted with 1,5 ml hexane and 1,5 ml of 5% w/v NaCl and combined with the first extraction. The organic phases were evaporated under nitrogen. FAMEs were analyzed by gas chromatography–mass spectrometry (GC/MS) at InMet (Rosario, Argentina). After resuspension in 0.5 mL of hexane, FAMEs were run in a GC TRACE 1300 Mainframe MS 230V coupled to a MS ISQ single quadrupole (Thermo Scientific, USA) with a capillary column TG-5MS (30 m, 0.25 mm i.d.; Thermo Scientific, USA). Operating conditions: column programmed from isothermal for 5 min at 140 °C, then 140 °C to 250 °C at 5 °C min−1, and isothermal for 1 min at 250 °C. Helium carrier gas flow was 70.5 ml min−1; split 1/45. Injector, ion source, and interface temperatures were 200 °C. The mass spectrum was continuously acquired from 40 to 600 m/z in full scan mode. The areas were determined by integrating SIM-74 ion.

### Flipper TR membrane tension staining, fluorescence lifetime imaging and analysis

After regular cell culture at 37°C (described above), cells were left at room temperature for 20 mins for temperature equilibration adaptation. Then, 0.75 uL of 1 mM Flipper-TR (Spirochorme, Switzerland) was added to 200ul of cell culture with an OD600∼0.5 and incubated for 7 mins. Then, 1 uL of stained bacterial culture was placed in a Matek dish and covered with a 1.5% agarose pad. To minimize fluorescence, the glass coverslip bottom of regular Matek dishes was replaced by clean 40 mm glass coverslips (Bioptechs Inc, USA). Glass coverslips were here cleaned by 1 hour of sonication in 1 M KOH after 3 rounds (1 hour) in fresh MQ water sonication.

Cells were imaged at room temperature in LSM 880 inverted confocal microscope (Zeiss, Germany). Using Zeiss software ZEN, a 20 MHz 440 nm pulsed laser was selected to image the cells. Scanning speed and zoom were chosen to achieve an imaging rate of 1 second per frame in continuous mode. Pinhole was set to 1 Airy Unit. Then, emitted light passed through a 550nm long pass filter and detected by a Time-Correlated Single Photon Counting (TCSPC) unit (Picoquant, Germany). SymPhoTime 64 software (Picoquant, Germany) was used to acquire data. Several cells in field of view (FOV) were imaged from 60-90 seconds (1fps) in order to be able to visually identify the lipid membranes compared to background and being able to manually select regions of interest accounting for at least 10^4 photons (Roffay et al., 2023). Laser power settings and imaging times were carefully chosen to minimize the photodamage.

Analysis was carried out in SymPhoTime 64 software (Picoquant, Germany). Due to cell size (width ∼0.7um) and our confocal z resolution, the FOV imaging comprised noise from the entire coverslip surface in the form of point-like noise in locations where no cells were found. Most probably, this signal corresponded to any kind of lipid aggregation (e.g. secreted lipid vesicles) or any hydrophobic aggregation from the media. Therefore, the determination of membrane tension lifetime for the entire FOV, comprising this noise, showed no consistent trend. It is worth to note that here used sample preparation methods were optimized to minimize the presence of this point-like noise. To solve this problem, we manually selected regions of interest (ROIs) corresponding to photons originating from mainly double membranes (primarily septa and adjacent cells) with the assistance of thresholding and magic-wand tools in SymPhoTime 64. Photons from different cells were analyzed per field-of-view and fitted to a two-exponential function. To obtain reliable fits, the number of photons must be ∼10^4^ in agreement with previous studies (Roffay et al., 2023). Due to a low membrane-photon count per cell, single-cell membrane tension profiles were technically infeasible. Therefore, an average membrane tension was reported as a function of membrane-excess, by integrating photon signal from multiple cells. Data points correspond to one Field of View (FOV) per independent sample. For each condition, we measured at least three 3 independent samples and >4 FOVs per sample.

### Laurdan lipid packing bulk assay

Laurdan experiments were performed using prior protocols for *B. subtilis* (Scheinpflug et al., 2016; Wenzel et al., 2018). Briefly, cells were grown to an OD600∼0.5. At room temperature, cells were filtered with a 10 µm filter to avoid chained cells. After filtration, cells were spun down for 1 min at 13.000×g in a bench centrifuge to remove media. Cell pellets were resuspended in prewarmed PBS+0.1% glucose to reach an OD∼0.5. 10uL of freshly prepared Laurdan (Invitrogen, USA) with a concentration of 1 mM in DMSO was added and incubated for 5 mins at 37°C. Then, cells were washed with prewarmed fresh PBS/glucose 3 times. After the last wash, resuspend to obtain an OD600∼0.5. Lastly, cells were placed on a roller drum grown at 37°C for 20 mins.

Laurdan fluorescence emission spectra were obtained in a Fluorolog fluorimeter (Horiba, Japan) using a Xenon-arc lamp as light source. Temperature was controlled by a circulating water bath and set to at 37°C. Sample homogeneity was maintained by continuous magnetic stirring. Laurdan was excited at 350 nm with an excitation bandpass of 2.5 nm width to minimize bleaching. Laurdan spectra were collected from 390-550 nm in 5 nm steps. The generalized polarization coefficient was calculated 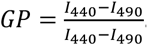. For hyperosmotic shocks experiments, 600 uL of cells at 37°C with OD600∼0.6 was mixed with prewarmed 300 uL fresh PBS or PBS with 1.5M / 3M sorbitol (final osmolarity of 0.5 M / 1M), immediately vortexed for few seconds, and placed in the fluorometer to start acquisition. Laurdan spectra as a function of time was recorded every 30 sec. At least 3 independent experiments were carried out per condition.

### Nile red staining and cell size calculation

1uL of fresh aliquots of Nile Red (5 mg/ml) was added to 1000 ul cells with OD600∼0.5 to account for a concentration of 5 ug/ml. Samples were excited with a 561 nm laser, in widefield illumination, using a 100X objective with NA=1.49 (Nikon, Japan) and a time exposure of 100ms. Emitted light passed through a 605 nm long pass filter and was detected by an Orca-Flash4.0 sCMOS camera (Hamamatsu, Japan) operated through NIS-Elements software (Nikon, Japan). Raw images were cropped by the user to generate ROIs suitable for batch analysis. ROIs mostly avoided clusters of cells to facilitate segmentation. Size determination was performed in batch mode by Morphometrics (Ursell et al., 2017) in a Matlab environment. Variables representing area, length and width were saved and used for plotting and further analysis.

### FtsZ-YFP-mts protein purification

FtsZ-YFP-mts was purified as described before (Osawa et al., 2008). Briefly, the protein was expressed from a pET-11b expression vector and transformed into *E. coli* strain BL21. Overexpression was performed at 20 °C. Cells were lysed by sonication and separated by centrifugation. Then, protein was precipitated from the supernatant, adding 30% ammonium sulfate and incubating the mixture for 20 min on ice (slow shaking). After centrifugation and resuspension of the pellet, the protein was purified by anion exchange chromatography using a 5 × 5-ml Hi-Trap Q-Sepharose column (GE Healthcare, 17515601). Protein purity was confirmed by SDS-PAGE.

### GUV preparation: electroswelling and double emulsion FtsZ encapsulation

Electro-swelled giant vesicles (GUVs) were made of different lipids (Avanti Lipids, United States) dissolved in chloroform. Using teflon chambers, three drops (∼1 µl) were carefully seeded in Pt wires and rapidly air-dried. After 1-h vacuum for further chloroform drying, GUVs were swelled in 250-mOsm sucrose solution at 10 Hz for 2 h and 2 Hz for 1 additional hour (detachment). For the case of *E. coli* and *E. coli*/Cholesterol 90:10, lipid concentration was 3 mg/ml. For EggPG/POPG 80:20, lipid concentration was 1 mg/ml. Flipper-TR (from DMSO solution) at 0.75% mol was added to the chloroform lipid mixture. GUVs are diluted (1:20) in buffer (125-mM KCl, 25-mM Tris-HCl, 2-mM MgCl2 pH 7.5)

FtsZ encapsulating vesicles were produced using droplet emulsion transfer (Pautot et al., 2003). Lipid composition was EggPC/DOPG 80:20 (Avanti, AL, United States) with 0.05% mol ATTO655-DOPE (ATTO-Tech GmbH, Germany). Briefly, vacuum-dried lipids were dissolved in mineral oil (Sigma, Germany) to reach a final concentration of 0.5 mg/ml. To form lipid vesicles, two interfaces are required: outer and inner interface. In a reaction tube (A), 500 µl of lipid + oil mixture was added to 500 µL of buffer (125-mM KCl, 25-mM Tris-HCl, 2-mM MgCl2 pH 7.5). At the oil–water interface, a lipid monolayer was assembled after 30 min. In a second reaction tube, 15 µl of protein master mix was added 500 µl of lipid + oil and vigorously vortexed for 2 min to obtain a homogenous cloudy emulsion. The inner monolayer was rapidly formed (∼2 min). This protein master mix was composed of buffer, 20% OptiPrep (Density Gradient Medium, Sigma, Germany), protein and GTP. FtsZ-YFP-mts and GTP final concentrations were 1.65 µM and 1.4 mM, respectively. Therefore, the emulsion was transferred to the reaction tube (A) and centrifuged at 100 g for 3 mins. Finally, the oil-based supernatant is discarded and 300-µl final vesicles are 1:2 or 1:3 diluted in fresh outer buffer.

Electroswelled vesicles or vesicles encapsulating FtsZ were imaged in home-made reaction chambers with passivated glass. Passivation was carried out by incubating 5 mg/ml b-casein for ∼3 hours on fresh plasma cleaned glass.

### Live cell imaging and FtsZ time evolution: ring thickness and constriction rates

Cells carrying native FtsA-mNeonGreen (bDR018, bDR112) were excited with a 488 nm laser, in widefield illumination, using a 100X objective with NA=1.49 and a time exposure of 200ms. Emitted light passed through a bandpass 500-550 nm filter. Images were acquired every 30 sec. For some acquisitions, the membrane (Nile Red) was simultaneously imaged with a rate of 1 image every 2.5 mins.

Time lapses for the mNeonGreen comprised at least 100 time points. ROIs from raw acquisition were manually selected to avoid cell clusters and facilitate ring detection. FtsZ rings with a 30-pixel diameter were detected using a Laplacian of Gaussian (LoG) detector in TrackMate (Tinevez et al., 2017). Quality thresholds were kept constant for different conditions and strains. Ring tracks were formed by TrackMate using “Linking max distance=20 pixels”, “Gap-closing max distance=20 pixels” and “Gap-closing max frame gap=0” parameters set.

A first Matlab script (tracking_rings_2.m) imported the tracking file from Trackmate together with the timelapses (pre-processed with a mean filter with radius=2 pixels) to generate timelapses per ring. Timelapses per ring were manually curated to discard wrongful tracks. Next, a Fiji/ImageJ script (process_Folder_zproj.ijm), performed a temporal average intensity projection per ring track in a batch mode. A second Matlab script (zcond_constr_final.m) showed the temporal average intensity projection and allowed the user to define two straight lines: one overlapping with the ring (“division plane”) and another perpendicular defined as “centerline”. Next, over the trajectory defined as “division plane”, the script calculated and displayed the corresponding kymograph, from the timelapse per ring, and allowed the user to draw two straight lines (constriction rates) if the quality of the kymograph is accepted by the user. On the other hand, over the trajectory defined as “centerline”, the script fitted the intensity profile to a gaussian function and defined a threshold value (thr) to determine which pixels are considered ring: thr=min_p+(max_p-min_p)/2 with the minimal value min_p and max_p the maximal value of the intensity profile. Using this threshold, ring thickness and the brightness per ring (with background subtraction) were computed. Finally, a third Matlab script (analysis_cond.m) filtered data using R2 >0.85 from the gaussian fit and constriction rates <70 nm/min.

### Total internal reflection microscopy (TIRF) for FtsZ treadmilling and analysis

Cells carrying native FtsA-mNeonGreen (bAB167, bDR018) were excited with a 488 nm laser, in TIRF mode, using a 100X objective with NA=1.49 and exposure time of 1s. With this exposure time, laser power was chosen to minimize photodamage yet obtain a good signal to noise. Emitted light passed through a bandpass 500-550 nm filter. 120 time points with an imaging rate of 1fps were acquired. ROIs from raw acquisition were manually selected to avoid cell clusters and facilitate single molecule -like-events or “treadmilling events’’. These events, with an estimated diameter of 6 pixels, were detected using a Laplacian of Gaussian (LoG) detector in TrackMate. Tracks were formed by TrackMate using “Linking max distance=6 pixels”, “Gap-closing max distance=6 pixels” and “Gap-closing max frame gap=0” parameters set.

A Matlab script (analysis.m) imported the tracking file from Trackmate together with the timelapses (pre-processed with a mean filter with radius=2 pixels). The script loaded every trajectory and calculated the track time length, the linearity of the trajectory (R2 from a linear regression) and track displacement. With these parameters, the scripts attempted to emulate manual kymograph slope determination in order to compare to manual previous measurements (Bisson-Filho et al., 2017; Squyres et al., 2021). Therefore, treadmilling tracks were formed by selecting linear trajectories (R2>0.8) with respective constant velocity (R2>0.8), tracks with time length between 7 and 50 seconds, and a total displacement between 6 and 10 pixels. In addition, for each selected trajectory, the script saved an image of 60x60 pixels, centered in the trajectory, showing a superposition of the first image and the trajectory itself. In this way, images corresponding to non-perpendicular trajectories were inspected and deleted. A last Matlab script (rectificar.m), discarded the velocity of deleted images as well as velocities below 10 nm/sec and above 60 nm/sec.

### 2D-SIM for Nile Red Imaging

Samples stained with Nile Red (described above) were excited with a 561 nm laser, in a 2D-SIM mode from a NSIM Nikon Microscope, using a 100X objective with NA=1.49 (Nikon, Japan). Emitted light passed through a 605 nm long pass filter and detected by an Orca-Flash4.0 sCMOS camera (Hamamatsu, Japan) operated through NIS-Elements software (Nikon, Japan). In the 2D-SIM mode, 9 images corresponding to 9 rotations in the illumination profile were acquired with an exposure time of 200 ms. Images were reconstructed in NIS-Elements software (Nikon, Japan) having “Illumination Modulation Constrast”=“auto” and “High Resolution Noise Suppression”= “1”. Reconstructed images were used in Figs.1C, S1B, and S1D.

### TEM of septal thickness

Strain PY79 and bDR110 were cultured as mentioned above. To achieve xylose dilution, liquid culture of strain bDR110 was diluted into fresh CH media (1:25 dilution) with different IPTG concentrations in flasks on a shaker at 185 rpm, 37°C. After 5.5 hours of depletion, cells were collected by centrifuge at 3000 rpm, for 3 min. Aliquots of cell pellets were transferred to the 0.2-mm deep well of aluminum planchettes (Technotrade International, Manchester, NH) and high-pressure-frozen in a BAL-TEC (Oerlikon, Balzers, Liechtenstein) HPM-010 High-Pressure Freezer. The frozen samples were freeze-substituted in 0.25% glutaraldehyde/0.1% uranyl acetate in acetone at −80°C for 3 days, then gradually warmed to −20°C, infiltrated with Lowicryl HM20 embedding resin (EMS, Hatfield, PA), and finally polymerized in BEEM capsules under UV illumination at −45°C. Sections with a nominal thickness of 60 nm were collected on formvar-coated copper or nickel EM grids. Images were taken by conventional transmission electron microscope JEOL 1200EX-80kv equipped with an AMT 2k CCD camera. Septum thickness was measured by FIJI software.

### Strain Construction

DNA fragments were assembled by polymerase chain reaction amplification and Gibson cloning. Gibson products were transformed directly into competent *B. subtilis* PY79 and integrated directly into the chromosome by homologous recombination with homology regions that were engineered at each end of the construct. Each construct was initially transformed into the WT background (PY79). Transformants were selected by growth on LB plates containing the appropriate antibiotic. The resulting strains were verified by polymerase chain reaction and, when appropriate, by sequencing.

**bDR018**: (amyE::tet-pXyl-accDA + ftsA::ftsA-mNeonGreen(SW)) was generated by transforming bAB167 (*ftsA::ftsA-mNeonGreen-SW*) with gDNA from bSW305 (*amyE::tet-pXyl-accDA)*

**bDR026**: (ycgO::spec-LacI-pyHyperSpank-accDA) was generated ligating 4 fragments with Gibson assembly: i) ycgO-spec (oMD247-oJM28), ii)LacI-pHyperSpank (oMD234-oWM232), iii) accDA(oDR031-oDR032), iv) ygcO-down (oMD257-oMD252).

**bDR125**: (ycgO::spec-LacI-pyHyperSpank-accDA & ftsA::ftsA-mNeonGreen(SW)) was created by transforming bAB167 (Native ftsA-mNeonGreen SW) using gDNA from *ycgO::spec-LacI-pyHyperSpank-accDA* from bDR026.

**bDR110**: (*ycgO::spec-LacI-pyHyperSpank-accDA* + *ΔsepF:tet + ezrA::cat-pXyl-ezrA*) was created in 5 steps:

1. Py79 was transformed with *ycgO::spec-LacI-pyHyperSpank-accDA*, which was created by ligating 4 fragments with Gibson assembly: i) ycgO-spec (oMD247-oJM28), ii) LacI-pHyperSpank (oMD234-oWM232), iii) accDA(oDR031-oDR032), iv) ygcO-down (oMD257-oMD252).
2. *ΔsepF:tet* was created by ligating 3 fragments with Gibson assembly: i) homology-up (oMH43-oMH98), ii) tet cassette (oJM28-oJM29), iii) homology-down (oMH20-oMH21).
3. The *ycgO::spec-LacI-pyHyperSpank-accDA* strain was transformed with gDNA from Δ*sepF:tet*.
4. ezrA::cat-pXyl-ezrA: ligated 4 fragments with Gibson assembly: i) homology up (oMH53-oMH54), ii) cat cassette (oJM29-oJM28), iii) pXyl (oMD73-oMD226), iv) homology-down (oMH14-oMH56).
5. The ycgO::spec-LacI-pyHyperSpank-accDA, Δ*sepF:tet* strain was then transformed with PCR amplified fragment *ezrA::cat-pXyl-ezrA* (oMH53-oMH56).

**bDR112:** (*ycgO::spec-LacI-pyHyperSpank-accDA*, *ΔsepF:tet, ezrA::cat-pXyl-ezrA*, ftsA::ftsA-mNeonGreen(SW)) was created by transforming bAB167 (Native ftsA-mNeonGreen SW) was transformed gDNA from *ycgO::spec-LacI-pyHyperSpank-accDA* from bDR110. It was then transformed with *ΔsepF:tet* DNA, and then ezrA::cat-pXyl-ezrA PCR products from bDR110.

## SUPPLEMENTAL MATERIAL

### SUPPLEMENTAL FIGURES

**Figure S1.**
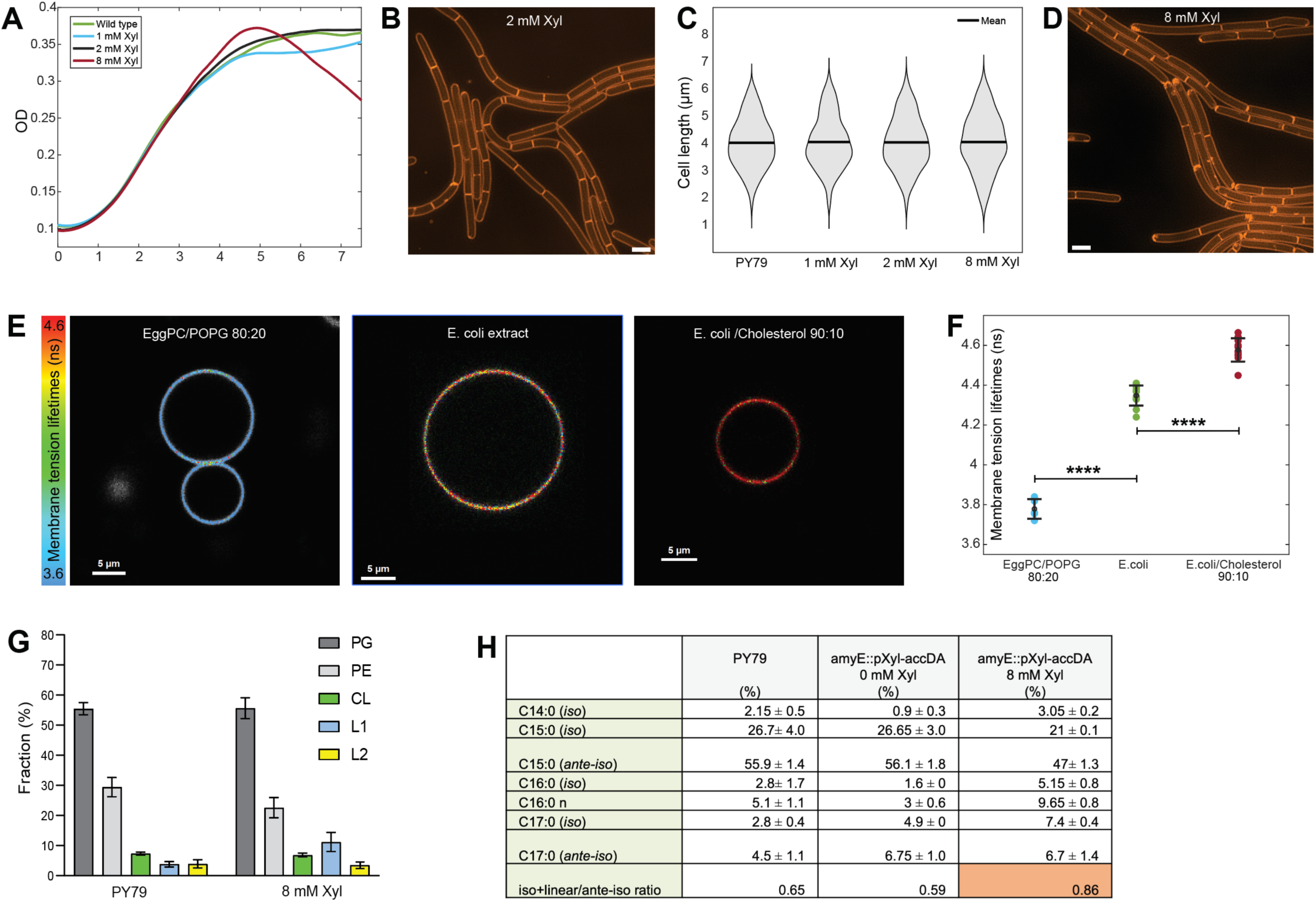
**A.** bSW305 growth in CH media as a function of different *accDA* expression levels (xylose) assayed by OD600 readings of liquid culture. **B.** 2D-Structured Illumination Microscopy (2D-SIM) images of bSW305 cells stained with Nile red with *accDA* induced with 2 mM xylose. Scale bar is 1 µm. **C.** Cell length distribution for different xylose inductions of *accDA* in bSW305. Scale bar is 1 µm. **D.** 2D-SIM images of bSW305 cells stained with Nile red with *accDA* induced with 8 mM xylose. Scale bar is 1 µm. **E.** Fluorescence lifetime microscopy images of GUVs made with different lipid compositions stained with Flipper-TR dye. **F.** Plot of FlipperTR lifetimes for different GUVs with different lipid compositions shown in **E.** Each data point corresponds to the fitted value gained from all GUVs within one field of view. P-values are in Table S1. **G.** Phospholipid polar head group distribution for bSW305 with 8 mM xylose accDA levels compared to PY79 wildtype wherein PG represent phosphatidylglycerol, PE:Phosphatidylethanolamine, CL: cardiolipin and L1&L2 unidentified polar elements in the TLC analysis. **H.** Fatty acid distribution of PY79 (wild type cells) and bSW305 with 0 mM and 8 mM xylose. AccDA levels were measured by mass spectrometry.

**Figure S2.**
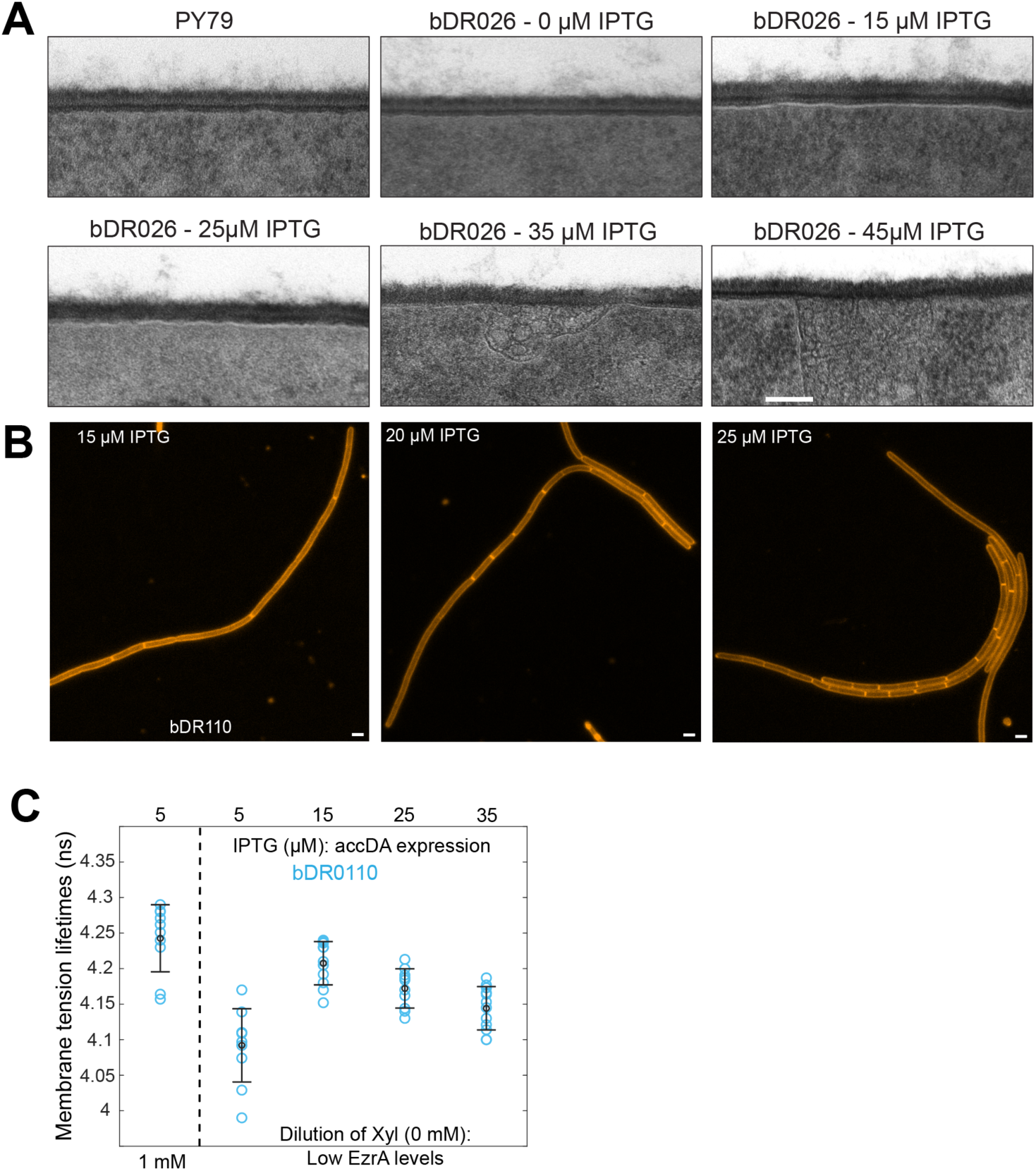
**A.** Transmission Electron microscopy images of bDR026 with *accDA* induced with different amounts of IPTG. Scale bar = 100 nm. **B.** Widefield fluorescence images of bDR112 cells stained with Nile red for different IPTG inductions of *accDA* in EzrA depleted *ΔsepF* cells. Scale bar = 1 µm. **C.** Plot of FlipperTR lifetimes at different for bDR110 at different IPTG inductions of *accDA* at high EzrA (1 mM xylose) and after EzrA was depleted for 4 hours. Photons from different single cells were analyzed. Each data point corresponds to the fitted value gained from all the cells within one field of view.

**Figure S3.**
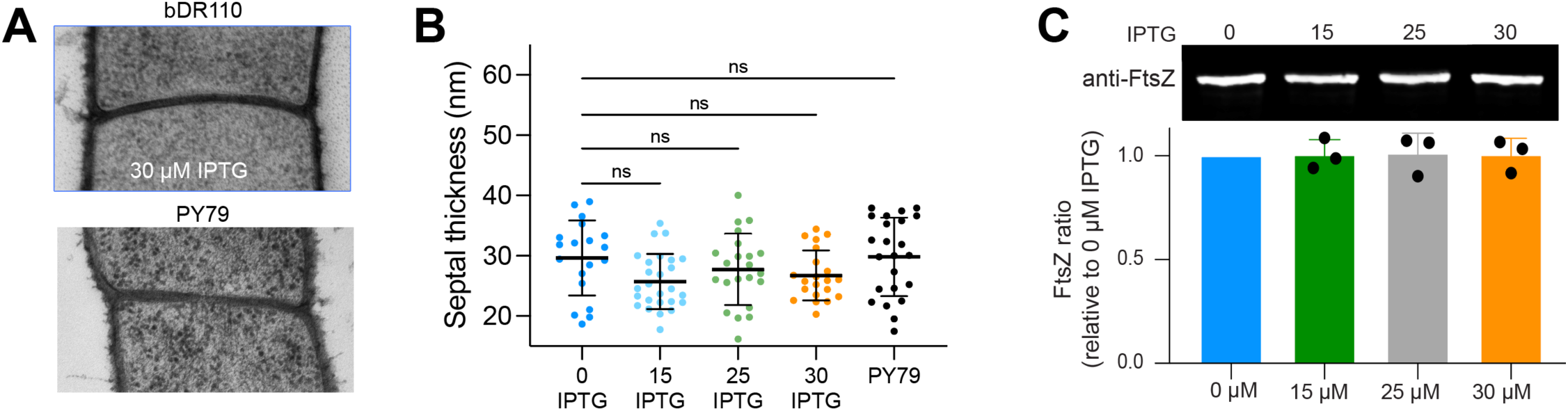
**A.** *Top* - EM images of septa from EzrA depleted bDR110 cells induced with 30 μM IPTG. *Bottom* - EM images of septa from wild-type cells. **B.** Septum thickness for bDR110 for different *accDA* inductions with 0mM xylose compared to wild type (PY79). Error bars are SD. P-values are in Table S1. **C.** Immunoblots of FtsZ in strain bDR110 with *accDA* induced at 0 µM, 15 µM, 25 µM, and 30 µM IPTG. Quantification of relative FtsZ level to strain bDR110 induced with 0 µM IPTG. Bar represents mean ± SD, P> 0.997, ns, ordinary one-way ANOVA.

**Figure S4.**
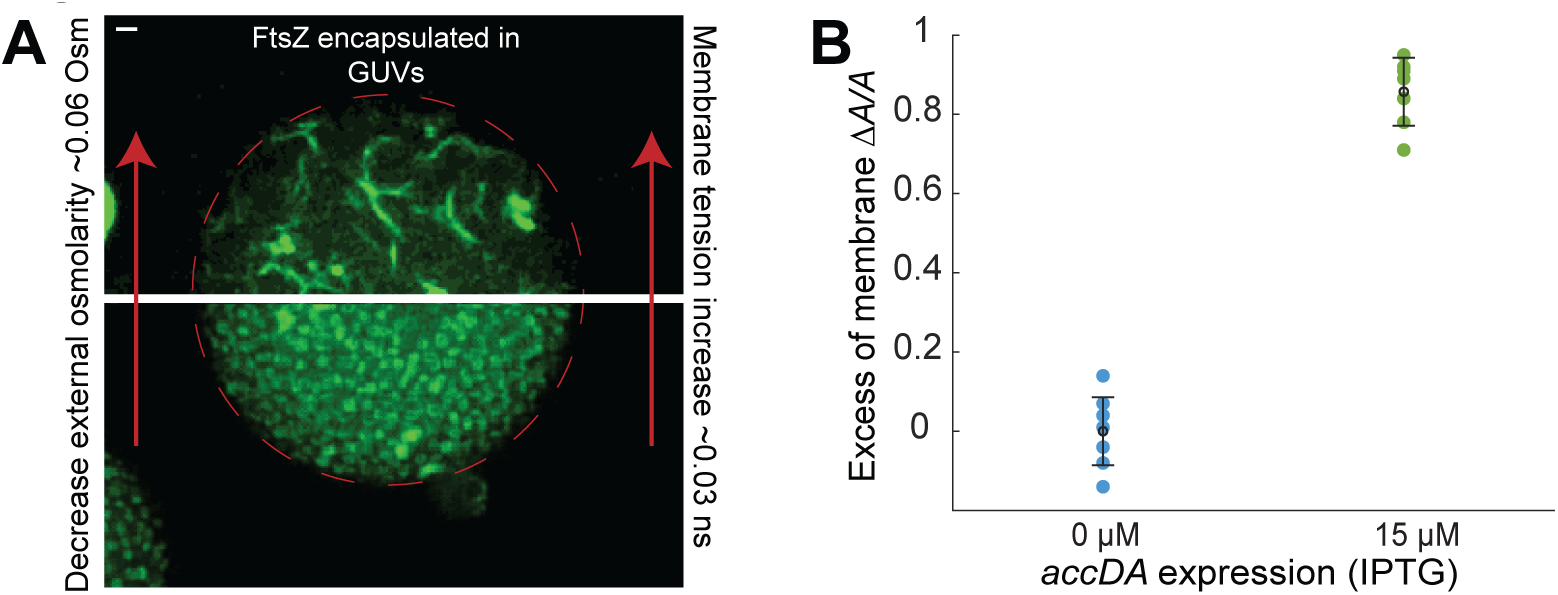
**A.** FtsZ-YFP-mts form rings with high curvature inside GUVs with low membrane tension. When membrane tension is slowly increased by ∼0.03 ns, rings turn into longer bundles with a smaller curvature. Scale bar is 1 µm. **B.** *De novo* total lipid synthesis was quantitated by measuring the incorporation of [^14^C] acetate into bDR026 with *accDA* induced at 0 μM and 15 μM IPTG.

**Table S1.** Statistics.

| Figure | Comparison | P-Value | Method |
| --- | --- | --- | --- |
| 1B | De novo lipid synthesis 0 – 1mM Xylose. | <0.0001 | One way ANOVA |
| 1B | De novo lipid synthesis. 1 – 8mM Xylose | <0.0001 | One way ANOVA |
| 1B | De novo lipid synthesis. 0 – 8mM Xylose | <0.0001 | One way ANOVA |
| 1B | Cell size 0 – 1mM Xylose | 0.0203 | One way ANOVA |
| 1B | Cell size 1 – 8mM Xylose | 0.0314 | One way ANOVA |
| 1E | PY79 vs 1 mM Xylose | 0.0187 | One way ANOVA |
| 1E | PY79 vs 2 mM Xylose | <0.0001 | One way ANOVA |
| 1E | 1 mM vs 2 mM Xylose | 0.0030 | One way ANOVA |
| 1E | 2 mM vs 8 mM Xylose | 0.9456 | One way ANOVA |
| 2F | 5 mM vs 15 mM IPTG | 0.0414 | One way ANOVA |
| 2F | 5 mM vs 25 mM IPTG | <0.0001 | One way ANOVA |
| 2F | 15mM vs 25 mM IPTG | 0.0358 | One way ANOVA |
| 2F | 15mM vs 35 mM IPTG | 0.0001 | One way ANOVA |
| 2F | 25mM vs 35 mM IPTG | 0.2799 | One way ANOVA |
| 3E | Treadmilling – PY79 vs 30mM | 0.2129 | Mann-Whitney |
| 3E | Constriction rate - PY79 vs 30mM | <0.0001 | Mann-Whitney |
| S1F | EggPC/POPG 80/20 vs. E.coli | <0.0001 | One way ANOVA |
| S1F | EggPC/POPG 80/20 vs. E.coli+10%Cholesterol | <0.0001 | One way ANOVA |
| S3B | 0 IPTG vs. 15 IPTG | 0.0684 | One way ANOVA |
| S3B | 0 IPTG vs. 25 IPTG | 0.6274 | One way ANOVA |
| S3B | 0 IPTG vs. 30 IPTG | 0.2785 | One way ANOVA |
| S3B | 0 IPTG vs. PY79 | 0.9998 | One way ANOVA |

**Table S2.**
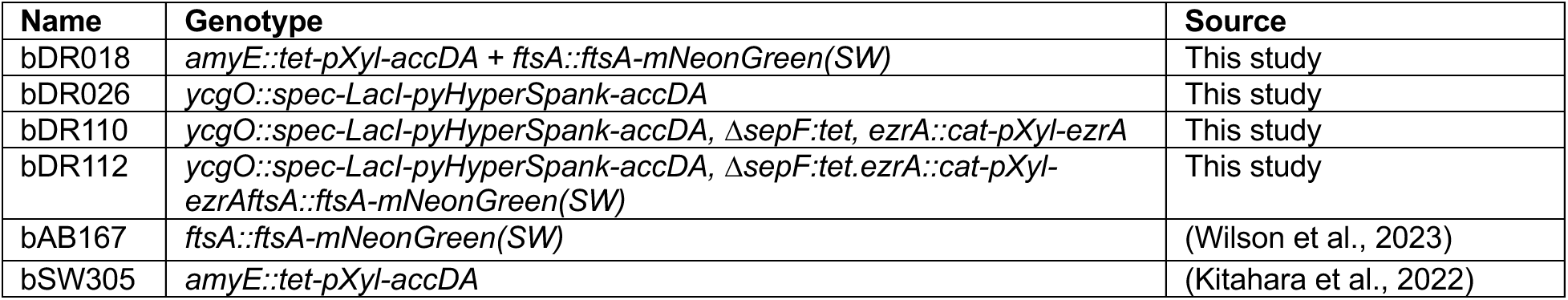
Strains Used in this Study.

**Table S3.**
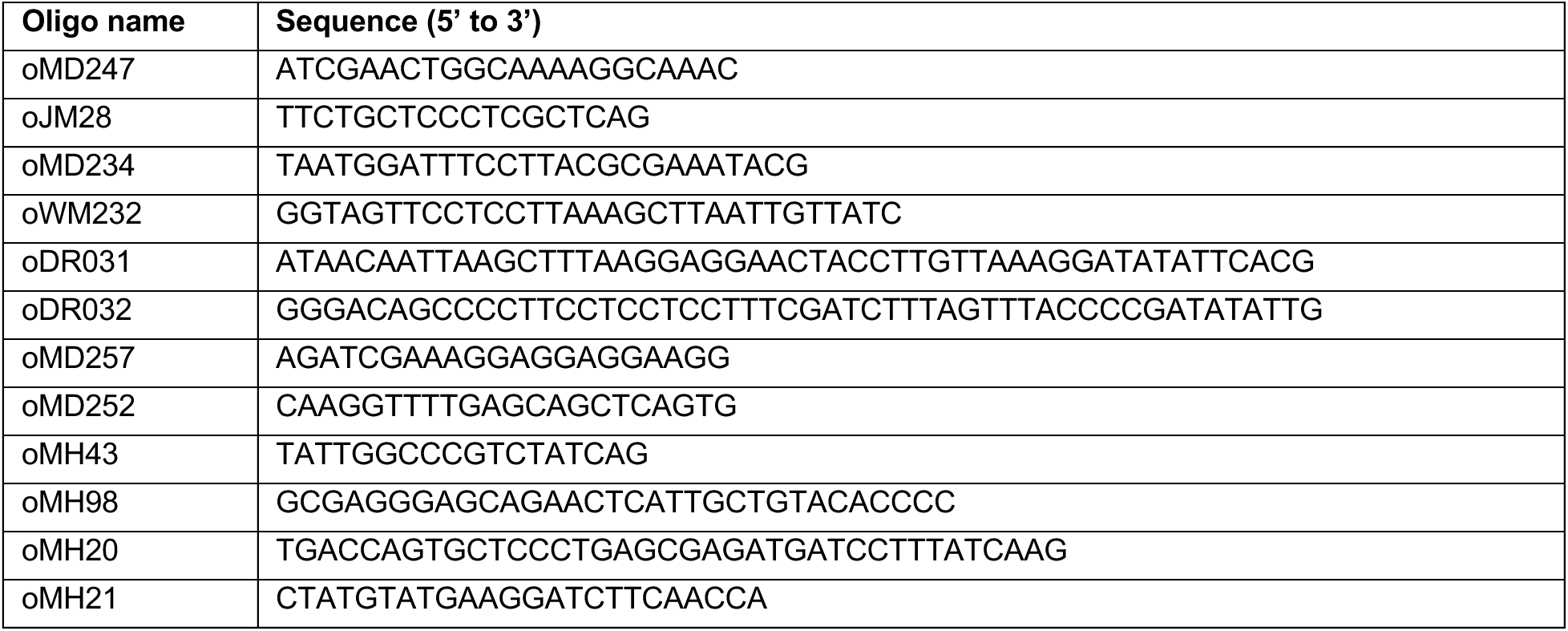
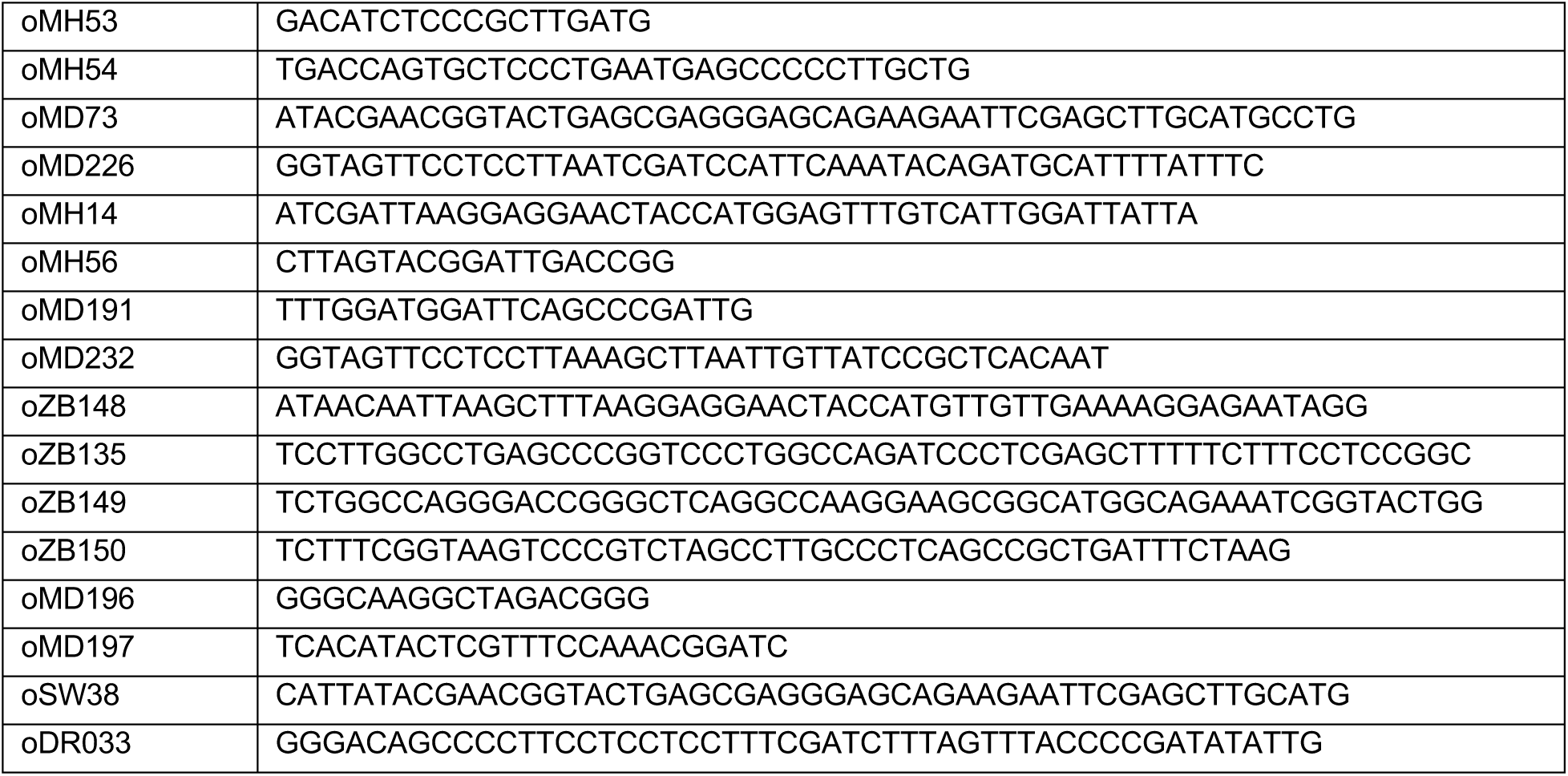
Primers Used in this Study.

## Supporting Information S1

In a cellular context, the membrane tension σ regulates how the membrane reacts to a local perturbation. The mechanical response to this perturbation (e.g., FtsZ ring along the circumference at midcell) occurs by the interplay of membrane stretching and flowing (Cohen and Shi, 2020; Djakbarova et al., 2021), in which flow represents the relaxation response of the membrane being locally stretched. This interplay of stretch and flow causes the membrane tension to propagate following a wave motion (Shi et al., 2018). Mathematically, the velocity of this flux moving towards a stress point is proportional to the membrane tension gradient Δ_σ_. By considering a bacterial cell of length *L*,

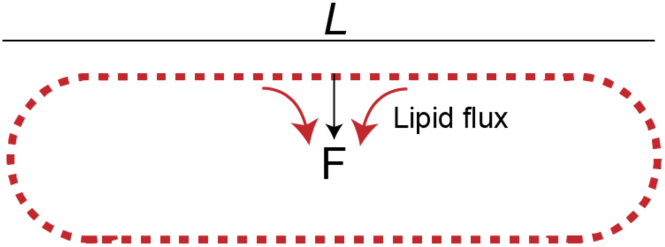

the velocity of lipid flux (*v*) towards the point of stress (*L*/2) is determined (Shi et al., 2022, 2018) by:

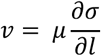

with 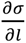 being the membrane tension gradient across the cell length and *μ* being the membrane mobility coefficient (Cohen and Shi, 2020). Here, we postulate that σ can be expressed as the sum of two components: the first associated with the magnitude of the local perturbation (*F*) and the other being related to a resistive component as a function of excess of membrane. The larger the excess membrane, the lower the resistance to flow and vice versa. Similarly, the larger the magnitude of the perturbation, the greater the membrane tension, causing a faster lipid flux. Then, this flux can be written as

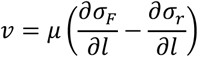

Since we measured the membrane tension σ*_m_* as a function of accDA levels, here we claim that σ*_r_* = σ*_m_*, implying that higher accDA levels correlate with lower resistance to flow and the opposite. By considering the resistant membrane tension σ_0_ as the highest expression level of accDA with no cell division (*v* = 0), then:

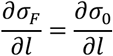

In view of this, the increase in the velocity of lipids being incorporated into the division site (*v*), as a function of accDA levels, can be expressed as

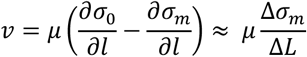

where Δ*L* represents the cell length difference as a function of accDA expression with respect to the highest expression level of accDA with no cell division: Δ*L* = *l_m_* − *l*_0_.

### Sequences of Fluorescent Proteins, Tags and Linkers

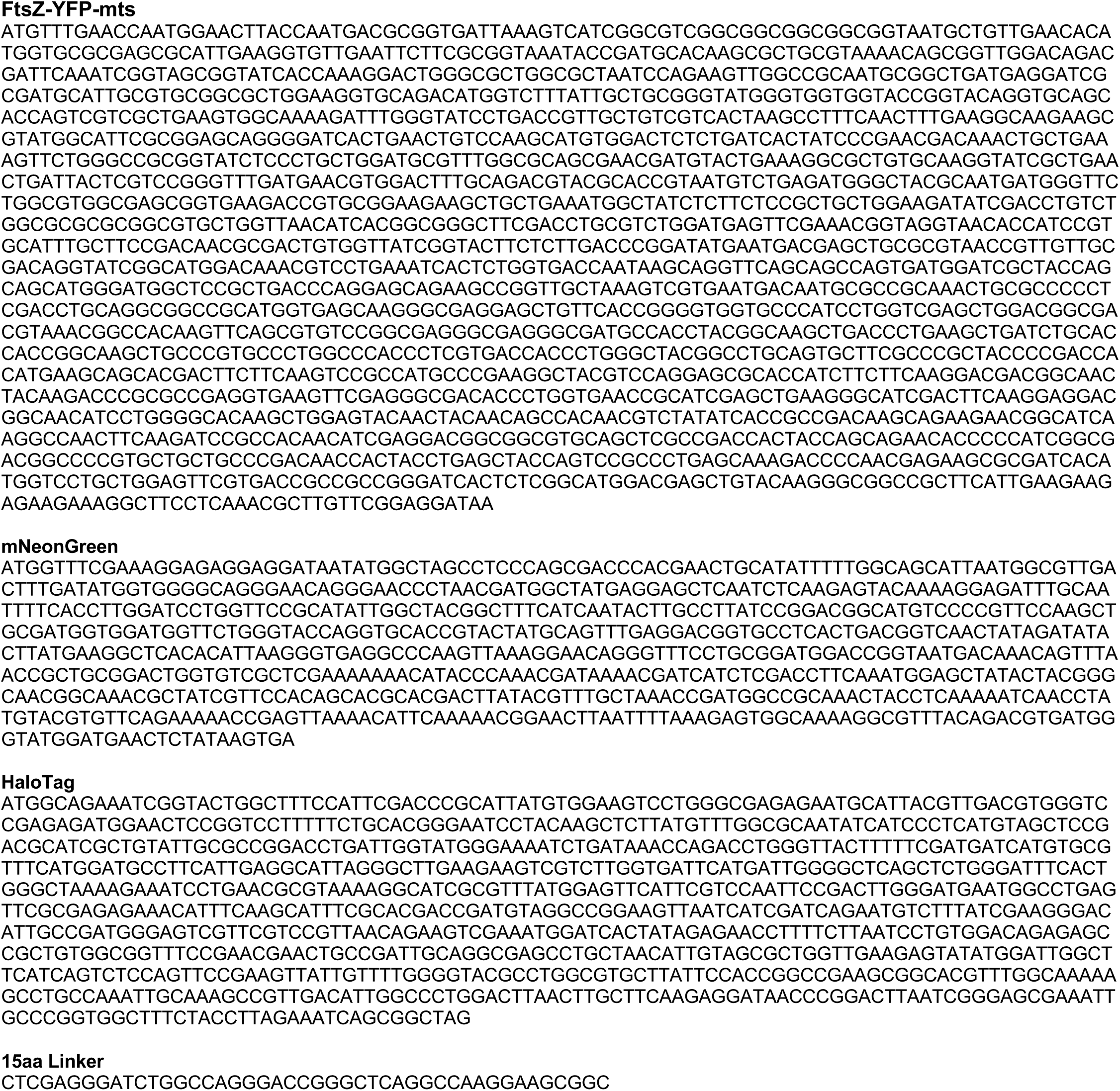

## Notes

### Competing Interest Statement

The authors have declared no competing interest.

